# Improvement in model flexibility reveals a minimal signaling pathway that explains T cell responses to pulsatile stimuli

**DOI:** 10.1101/2025.08.28.672930

**Authors:** Jun Allard

**Affiliations:** Department of Mathematics, Department of Physics & Astronomy, Center for Complex Biological Systems, University of California Irvine

## Abstract

Efforts to develop quantitative models must face a trade-off between interpretability and quantitative accuracy, which often disfavors interpretability. Here we adopt an operational definition of interpretability, specifically that a model is described by an arrow diagram wherein each arrow corresponds to a positive effect or negative effect of one component upon a process, and fewer arrows is more interpretable than more arrows. We then develop a method to add flexibility — and thus accuracy in fitting data — to mathematical models by relaxing functional form assumptions, while constrained by the same arrow diagram and thus the same interpretability. We apply this method to the T cell, where quantitative models are particularly needed, in part because of ongoing efforts to engineer T cells as therapeutics. One avenue of experiments exposes T cells to pulsatile inputs and measures their frequency response, finding several nonlinear frequency responses: high-pass, low-pass, band-pass, and band-stop. Using our modeling approach with enhanced flexibility, we show that a simple signaling model quantitatively captures the frequency response of CD69 surface expression, one of the correlates of T cells activation, with accuracy within the experimental inter-replicate standard error of the mean. Specific qualitative behaviors map to specific parts of the arrow diagram: Band-pass behavior can be explained by refractory de-sensitizing circuit (we refer to this as “first-aid icing a wound”). Band-stop behavior can be explained by removal-inhibition (we refer to this as “roommate interrupts my studying”). Apparent low-pass emerges naturally when total stimulation time is constant. We test the model on independent experimental datasets from multiple labs. Taken together, our results demonstrate the ability to achieve both quantitative prediction and interpretability in understanding cellular dynamics. Simple models may at first appear incapable of explaining complex data, but might indeed be able to by adding this modest flexibility.

## Introduction

T cells accept input in the form of ligands, which activate receptors, to produce cellular outputs. For the T cell, a major input is antigenic peptide presented by MHC, which activates the T Cell Receptor, and the cellular outputs include an array of immunological function, like cytokine secretion and surface molecule expression (1, 2). Due to their role in disease and therapeutics, there is an acute need for quantitative, predictive models of whole-cell responses from input to output (3–6).

Much work has been done on understanding the T cell’s response to inputs of different molecular identities, e.g., ligands with different affinities or accessory ligands, or different density (1, 7). One alternative way to experimentally characterize a complex system – with rich history of success across engineering and biology (8, 9) – is by presenting the system with pulsatile inputs, and varying the frequency or on-time (*T*_on_) and off-time (*T*_off_), as shown in Figure 1A. Indeed, it is hypothesized that T cells may be able to integrate pMHC signals across different cells, which would imply T cell behavior may be regulated by pulses of antigen encounters on the timescales of minutes to hours (11). Pulsatile inputs have been presented to T cells using optogenetics (10, 11). Fascinatingly, researchers found all four basic categories depending on cell type and specific output observed, roughly speaking:

- High-pass, in which gaps between pulses reduces the output;
- Low-pass, in which gaps between pulses enhance the output, to an even higher response than no-gap stimulation;
- Band-stop, in which intermediate-duration gaps reduce the output (11);
- Band-pass, in which intermediate-duration gaps enhance the output.

**Figure 1.**
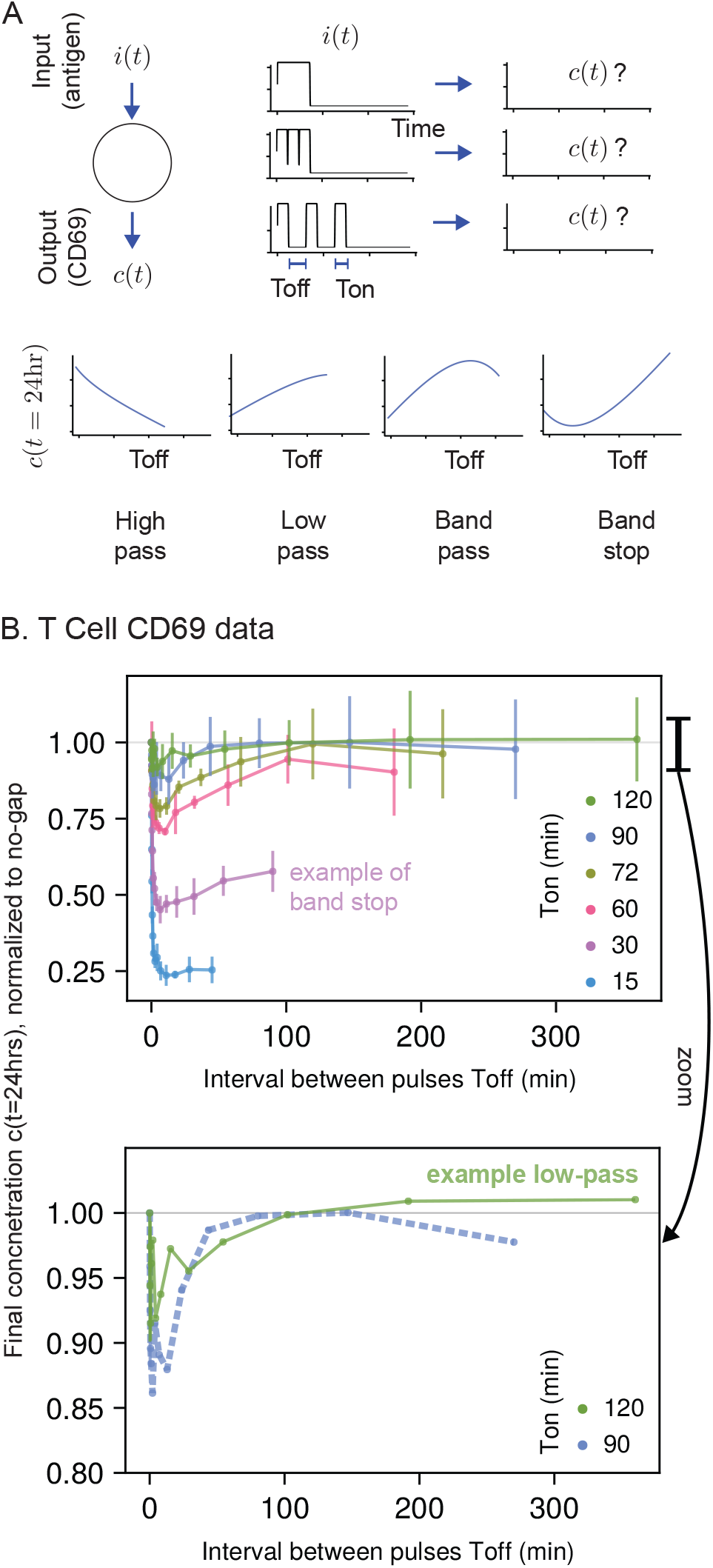
Experimentally-observed response of T cells to pulsatile stimuli reveals several categorically different features. A. Exposing a system to inputs *i*(*t*) with cycles of on-times *T*_on_ and off-times (intervals between stimulus, *T*_off_). In this case, the system is a T cell, and the input is optogenetically-activated antigen stimulation, and the output is surface expression of CD69. There are several responses, termed “frequency responses” and typically named after their response to frequencies (so, for example, “high pass” is the term for a strong response at high frequency, or short *T*_off_). B. The response of T cells to pulsatile exposure to antigen stimulation from (10). Surface expression of T cells was measured after 24 hours. Data is normalized to the response of a continuous 120 minutes of stimulation (*T*_off_ = 0). Error bars indicate the standard deviation between replicates (note that in figures with model fitting, the standard error of the mean is shown instead). “No-gap” refers to *T*_off_ = 0.

Several categorically different responses are evident in a dataset from (10) reproduced in Figure 1B, where the output is surface expression of CD69, a correlate of T cell activation.

One way to mathematically characterize a complex system is with phenotypic modeling (2), in which we seek a minimalist pathway, often expressed as system of coupled nonlinear differential equations that represent an arrow diagram, as in Figure 2A. Components in the minimalist pathway represent a coarse-grain view of molecular interactions — where each component contains complex biophysics and nonlinearities (12–18) and may not correspond to a specific molecule.

**Figure 2.**
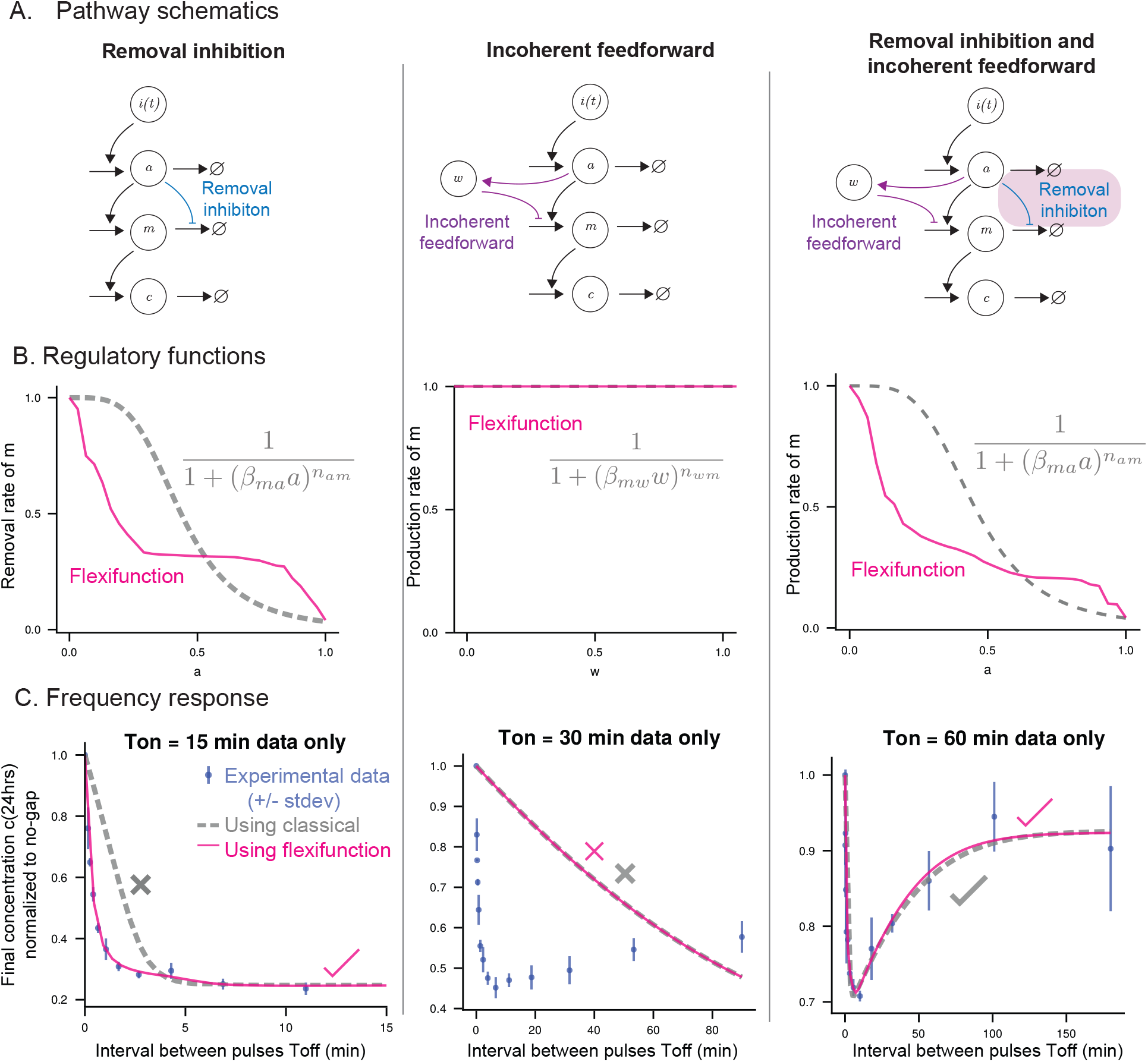
Model fitting to subsets of CD69 frequency response data with and without model flexibility. A Model pathways described as arrow diagrams, where each arrow is restricted to be either a positive relationship or negative relationship. The highlighted arrows represent one component regulating another, and either obey a mathematical expression or are allowed to explore a universal continuous positive function restricted to either increase or decrease. B The regulatory functions corresponding to highlighted arrows. Dashed line is classical model obeying equation. Solid line is the universal function, which we term a “flexifunction” for convenience. C Frequency response with single on-time series for both classical and flexible regulator functions. There are 6 data subsets, 4 classical models and 5 fleximodels, so there are 24 classical models+dataset pairs and 30 fleximodel+dataset pairs. Representative examples are shown of for 3 on-times, and a corresponding classical and fleximodel fit (dashed and solid line, respectively). Error bars indicate between-replicate standard deviation of data from (10). Checkmark or X indicate whether the root-mean-square discrepancy between model and single on-time series is above or below the experimental standard error of the mean. To see the root-mean-square fit of every model with every on-time subset, see Supplemental Figure S2.

Here, we ask, what is the phenotypic model that can explain the response of T cells to pulsatile stimulation observed in CD69 measurements by (10)? We specifically explore two motifs: the removal inhibition motif (sometimes called derepression (19) since it is repression of a repressor) and the incoherent feed-forward motif. Both of these motifs appear across biology and have been identified or proposed specifically for T Cell activation (4, 20–27).

Quantitative predictive models battle between interpretability and predictive accuracy (28). Models with many degrees of freedom often represent data more accurately, but may be more difficult to interpret. There are several transformative approaches to defeat this trade-off (29–31): approaches that approximate both a simple dynamical law and the data (32); approaches that begin with a large list of symbolic forms for mathematical expressions and perform sparse regression to select from these (33); and approaches that allow flexibility in specific terms in the differential equation using universal functions (34–37).

Here we adopt the goal of capturing the data to within its experimental uncertainty, with an arrow diagram where each arrow is a monotonic-increasing or monotonic-decreasing function (represented by arrow-head or bar-head, respectively, in Figure 2A). In other words, herein, our operational definition of interpretability is an arrow diagram where each arrow describes a positive effect or negative effect on its target process, and fewer arrows is more interpretable than more arrows. This is related to the above-mentioned approaches to balance interpretability and accuracy. One crucial difference in our goal is that the arrow diagram is viewed as a hypothesis to test and potentially reject. And so, the additional model flexibility must only explore models and parameters within this hypothesis, for example removal inhibition or incoherent feed-forward.

Using this method, we show that a simple signaling model quantitatively captures the frequency response of CD69 surface expression, with accuracy within the experimental inter-replicate standard error of the mean. Simple models may at first appear incapable of explaining complex data, but might indeed be able to by relaxing functional form assumptions.

## Results

### Phenotypic models of CD69 frequency response with classical mass-action and Hill-type assumptions

We began by listing five pathway arrow diagrams of increasing complexity. The first model we study is a linear pathway with 3 components *a*(*t*), *m*(*t*), *c*(*t*), shown schematically in Supplemental Figure S1A. In this model, each component is produced in response to the previous component, with the first component responding to the input *i*(*t*) and the last component *c*(*t*) identified as surface expression of CD69. The second simplest model we study is motivated by the experimental protocols of (10) and (11), both of which use an optogenetic tool called LOVTRAP (38) to activate and deactivate ligation of the receptor. This tool has its own internal timescale, which is estimated to be approximately 20 seconds in (10) but has exhibited timescales of ~ 500s in other cellular systems (38). To represent this, we studied a 4-component linear pathway *i*(*t*) → *l*(*t*) → *a*(*t*) → *m*(*t*) → *c*(*t*). Neither of these models are able to match, even qualitatively, the complex frequency response in Figure 1B, as shown in Figure S1.

Following the failure of the two linear architecture models to capture the significant nonlinearities in the CD69 data, we explored three more models with increasing complexity starting from the linear pathway model, shown in Figure 2A.

In the next model, rather than promote the production of a signaling molecule, an upstream component inhibits its removal. Removal inhibition (RI, sometimes called derepression (19) since it is repression of a repressor) has been suggested for several components of the T Cell activation pathway (20–22). We explore RI as a general phenomenon, but it has been specifically proposed, for example, TCR activation induces the HSP90-CDC37 chaperone to inhibit the degradation of Lck, which is an activator of the downstream signal (20). Calcineurin both activates a part of the pathway, NFAT, and dephosphorylates Lck at Ser59, which is an inhibitory phosphorylation (21). TCR activation also induces (indirectly, through Phosphoprotein Associated with Glycosphingolipid) the delocalization of Csk, which is an Lck inhibitor (22–24). In those cases, there is a component of the T Cell activation pathway that is both promoted, and also its inhibition is inhibited. So, in the RI model, *a*(*t*) inhibits the removal of *m*(*t*).

Then, we explore the incoherent feed-forward loop (IFF). In the incoherent feed-forward model, a new component *w* acts as a refractory variable, accumulating in response to *a* and reducing the accumulation of *m*. The IFF motif appears across biology (26, 39), is present in several mathematical models of T Cell activation (25, 40), was shown to be present in a minimal motif to explain T cell activation (4), recurs in complex adaptive systems (26), and is known to be sufficient for band-pass filtering (27). Finally, we explore a model that has both removal inhibition and incoherent feed-forward (RI+IFF).

One strategy to represent regulation, which has proven valuable over decades of mathematical biology research (6, 26, 41–44), is to use specific mathematical forms for the regulators containing 1-2 parameters (see equation in Figure 2B and Equations 9-16), and using these as terms in a coupled system of nonlinear ordinary differential equations. These functional forms can be derived from molecular interactions assuming mass-action kinetics plus, for example, multi-step irreversible processes (45) or intermediate-complex saturation (30, 46), yielding Hill-type equations as shown in Figure 2B. Graphs of these regulatory functions are shown in Figure 2B as dashed lines. We translate the arrow diagrams into models using these functional forms to represent the regulatory terms, and fit the parameters in these mathematical models to experimental data. Full equations and fitting algorithm are in Methods.

Before fitting to the full CD69 dataset, in this section we fit each model to a subset of the data corresponding to a specific on-time, out of the 6 on-times in the full dataset. For each model and on-time data series, we measure the root-mean-square error (rms) and report these in Supplemental Figure S2 and Supplemental Figure S3. Examples are shown in the 3 columns of Figure 2C (dashed lines). Two of the individual on time series can be fit by these models to within the standard error of the mean (a representative is shown in Figure 2C right, dashed line), and 9 can be fit within the standard deviation of the data. Most (15 out of 24) of the models and on-time series cannot be fit (representatives shown in Figure 2C left and middle).

### Subsets of CD69 frequency response data can be explained by simple models, when flexibility is added to model assumptions

The partial failure of the above models led us to think, *in vivo*, the individual arrows in the phenotypic models of Figure 2 may not obey simple functional forms. Many biophysical phenomena (surface-confinement, mechanics, phase separation) imbue nonlinearity into individual molecular steps (12–18), and anyway the nodes in the arrow diagrams are coarse-grain representations of many molecular interactions. So, we developed a modeling approach in which the regulatory functions are represented as arbitrary functions with determined monotonicity, in this case, always decreasing as a function of their input, which we refer to as “flexifunctions” for convenience. Note crucially that these models with flexifunctions correspond to the same arrow diagram as the classical models described above. For mathematical description and learning algorithm, see Methods. Briefly, the goal of the flexifunction approach is to describe a particular arrow diagram model, without making assumptions about the functional form of the arrows, but with assumption of the “sign” of the arrow, meaning whether it is a promotion/enhancement or inhibition/repression (so it can faithfully describe, e.g., RI or IFFL). Mathematically, this requires us to be able to learn from the set of all monotonically increasing or decreasing functions. There are several canonical ways to describe large families of functions, mathematically referred to as a basis. Canonical options include Chebyshev bases, Fourier series, multilayer perceptrons, cubic approximations, and piecewise-linear functions. Mathematically, as the number of degrees of freedom increases, any choice of basis will yield the same result, although computational implementation (including imperfections in the learning algorithm) may lead to differences in practice. We want sufficient degrees of freedom to capture the data, but not too much as to risk overfitting (see below and Supplemental Figure S5). We chose piecewise-linear functions for this project for their ease of implementation. However, we show in new Supplemental Figure S5 that we can also use cubic splines, which are smooth, and find similar results. From Supplemental Figure S5, we conclude that the choice of piecewise-linear basis is not the driver of the goodness of fit or gross model behavior.

The rms performance of these fleximodels is shown in Supplemental Figure S2. Since RI+IFF has two regulatory functions, either one or both could be represented as a flexifunction. Thus, there are 5 models × 6 data subsets = 30 fits. With this modest increase in modeling flexibility, 20 out of 30 agreed within the standard error of the mean, and 24 out of 30 were within the standard deviation. Representative examples are shown in the columns of Figure 2C (solid lines). The left and right examples agree with the data, and the middle example fails to fit the middle data.

One potential concern is that adding flexibility to the model could lead to over-fitting, in which the model learns to capture experimental error in the specific training set, and would not generalize well to repeats or new data. The monotonic flexifunctions we employ are approximators of universal monotonic function (subject to the constraints described in Methods), and their level of approximation is described by a parameter that controls the degrees of freedom in each flexifunction, *n*_dofs_ (defined in Methods). As the degrees of freedom *n*_dofs_ increase, the flexifunction approaches the universal function. This allows us to study the possibility of over-fitting. We perform cross-validation with increasing flexibility in the flexifunctions, shown in Supplemental Figure S4 and discussed in Methods. As expected, increasing flexifunction flexibility, via degrees of freedom, at first improves performance, but after around 32 or 64 degrees of freedom per flexi-function, the performance is maintained on the training data but lost on the testing data, thus suggesting an optimal flexibility to avoid overfitting. We note two further lines of evidence suggesting that the flexibility is not leading to overfitting: First, many fleximodels failed to fit on-time series data (e.g., Figure 2C). Second, the performance in Supplemental Figure S4 plateaus at large degrees of freedom, even on the training data, suggesting that there are structural constraints in the model preventing it from perfectly fitting the data. Both of those suggest the models are still sufficiently constrained to be used to reject hypotheses about the biological system.

### RI+IFF arrow diagram model can quantitatively explain full CD69 frequency response data, by relaxing assumption of functional forms

Having developed the flexible modeling method, we next sought to fit the model to the full dataset, learning the regulatory functions directly from the data if necessary. This process can be visualized in the table in Figure 3A: From left to right, model complexity increases (more arrows, more components). From top to bottom, model complexity is constant (same arrow diagram), but flexibility of individual terms is increased. The first model to agree with the full on-time, off-time dataset is the RI+IFF model with both regulatory functions represented as flexifunctions. The regulatory functions and frequency response are shown in Figure 3B and C (solid lines), along with the best-performing equivalent classical model (dashed lines). The model has 4 biophysical parameters representing timescales of the 4 components, in addition to the flexifunctions. The parameters of the best-fit model are: *τ*_*a*_ = 2.45, *τ*_*m*_ = 1.0, *τ*_*w*_ = 129.4, *τ*_*c*_ = 10^6^, in minutes. The timescale of *τ*_*c*_ attained its upper-bound and is much larger than the experimental timescale, so effectively *τ*_*c*_ → ∞.

**Figure 3.**
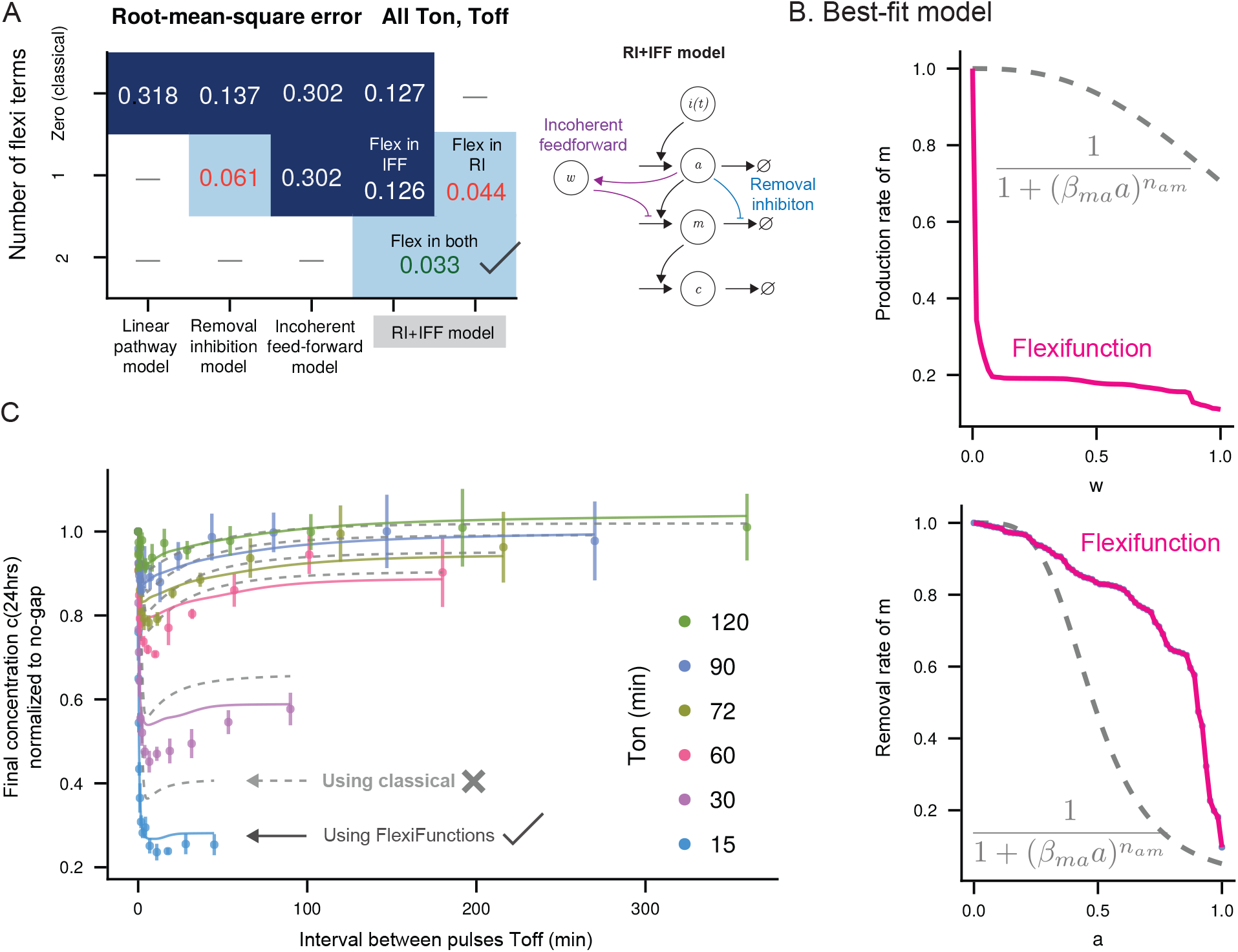
A simple model with removal-inhibition and incoherent feed-forward can fit full CD69 dataset, when flexibility is added to regulatory functions. A Best-fit root-mean-square error when fitting to all on-times and off-times simultaneously. Four arrow diagram models, with either zero (classical), 1 or 2 flexi-function terms. Models with best-fit rms below the standard error of the mean shown in green with a checkmark. Models with best-fit rms below the standard deviation shown in orange. B The best-fit model and the only model we explored that has root-mean-square error below the experimental standard error of the mean is the model with both removal-inhibition and incoherent feed-forward, and both of those associated regulatory functions were allowed to be flexible. Regulator functions of best-fit RI-IFF model (solid line) deviate significantly from the best-fit classical regulatory functions. (Note that the other parameters have also changed between models.) C Best-fit frequency response. Error bars indicate between-replicate standard deviation from (10). Solid lines are best-fit flexi model. Dashed lines are best-fit classical model.

Taken together, this demonstrates that a simple model may at first appear incapable of explaining complex data, but might indeed be able to do so, by relaxing functional form assumptions.

Inspection of the regulatory functions suggests an explanation for why classical model failed to fit. The data exhibits a dramatic decrease in response for even very short off-times, when on-times are short, and yet only moderate decrease for longer off-times or on-times. To capture this, the refractory inhibitor gives dramatic inhibition after a small build-up in *w*, and only moderately more inhibition as more *w* accumulates. We include the full time-series for all on-times and off-times in **Supplemental Movie 1**. A notable feature is that the first pulse leads to much larger *m* than subsequent pulses, when on-time and off-time are small. This is consistent with the regulatory function and the data.

The regulatory functions discovered by the learning function, shown in Figure 3B, are approximated by piecewise-linear functions, which appear slightly jagged. It might be beneficial to have a model in which the regulatory function is assumed to be smooth. This could either be for aesthetic reasons or due to prior expectations of the response of a cell (although note that all experimental measurements of intermediate input-output relationships appear rough, for example ppERK or NFAT in (10).) We can replace the piecewise-linear flexi function with a kernel-smoothed version, as shown in Figure S5A. The flexi-functions were smoothed using cubic approximators (47), with nonuniform knot placement (15 knots in [0, 0.2], 10 knots in [0.2, 1]) to increase resolution for low values of *w*. This has a minimal effect on the model behavior, as shown in Figure S5B. Note the model was not retrained, so this is a true test of model sensitivity, showing only weak dependence on the fine-grain details of the regulator functions.

While the best-fit line has a root-mean-square error below the average standard error of the mean of all data, it does not run through every data point’s error bars. Indeed, there are systematic mismatches, for example the steep dips in the data at low *T*_off_. It is possible that these could be included in the same arrow diagram by replacing the *m* → *c* term with a flexifunction, but we did not attempt this, to maintain our strict commitment to the original desired threshold of goodness of fit.

### Band-stop and low-pass arise in RI+IFF due to “roommate interrupts studying” and “icing a wound” phenomena

A widely-valued power of classical mathematical biology modeling is that their qualitative behaviors are often interpretable. We take a single on-time series that exhibits both band-stop (a local minimum in response versus *T*_off_) and low-pass (an enhanced response for larger *T*_off_, even above the no-gap stimulus), shown in Figure 4A.

**Figure 4.**
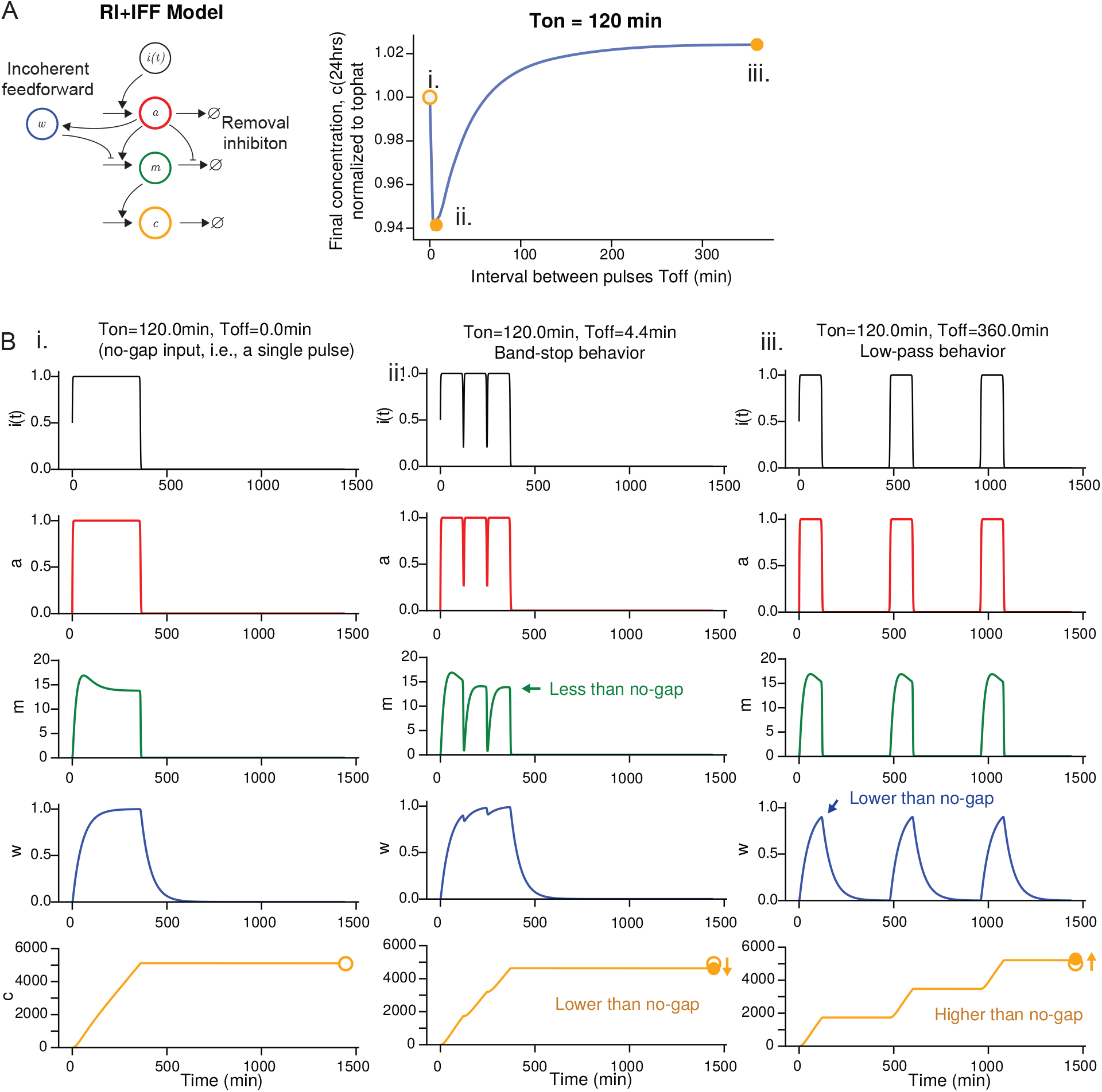
Qualitative features in frequency response have simple heuristic explanations. A Frequency response with single on-time series showing band-stop (the response is lower when there are intermediate gaps between pulses) and low-pass (the response is higher when there are large gaps between pulses). B Three selected time series, at off-times indicated by letters in (A). (i) tophat, i.e., one continuous pulse, (ii) intermediate gaps between pulses leading to lower response at 24 hours, (iii) long gaps between pulses leading to higher response at 24 hours.

First, band-stop behavior, or the decrease in response for moderate gaps between pulses. The *m* variable accumulates during stimulation, but decays rapidly when stimulus from *a* is removed due to its role in inhibiting removal. As a result, even short gaps in the stimulus prevent *m* from reaching its no-gap value (green arrow in Figure 4Bii). This harms the rate of increase in *c*. We refer to this as “roommate interrupts my studying” or “P.I. interrupts my coding” (48): in this analogy, the intermediate quantity *m* is a latent ability to focus and study or write code.

Next, low-pass behavior, or, the increase in response for large gaps between pulses, even higher response than no-gap stimulation. During no-gap stimulation, the refractory variable *w* accumulates gradually (blue curve in Figure 4Bi), limiting the steady rate of *c* accumulation. If there are gaps longer than the timescale of *w*, then *w* does not have a chance to impact *c* to its full effect (blue arrow in Figure 4Biii), leading to a stronger response. This is a known feature of biological systems that exhibit adaptation and self-regulation (49–52). One example is offered by blood flow to a wound after an injury: common first-aid advice is to apply ice to a wound, but to do so intermittently. In this analogy, the refractory variable *w* is your body’s homeostasis signal (compensatory vasodilation), which de-sensitizes you to the ice.

Note that both effects (band-stop arising from roommate-interrupts-my-studying and low-pass arising from adaptation) are features of the arrow diagram models, and arise in both the classical mathematically-expressed models and the models with flexifunctions.

### Apparent band-pass arises with constant total stimulation time, in all models

The experimental protocol of (10) used a constant total stimulation time in all measurements, for technical reasons (to control for consistent damage accumulation). And, as mentioned above, the response is measured at a fixed time (24 hours after the first pulse begins). In this protocol, it is possible that larger gaps between pulses (*T*_off_) give rise to stronger response — which we refer to as apparent band-pass — simply because there is less time between the last pulse and the measurement.

Indeed, even the linear pathway model can exhibit apparent band-pass under this stimulus protocol. This is shown in Figure 5A. Note that even though the linear pathway model can give apparent band-pass, it is still incapable of explaining the CD69 data, since the data exhibits both band-stop and band-pass in the same *T*_on_ data series, whereas the linear pathway model can only give apparent band-pass. To confirm that this is a feature of the protocol and not the model, we simulate the linear pathway model with a modified protocol where pulses continue until the measurement at 24 hours, in Figure 5C. Here, the model was incapable of increasing response strength.

**Figure 5.**
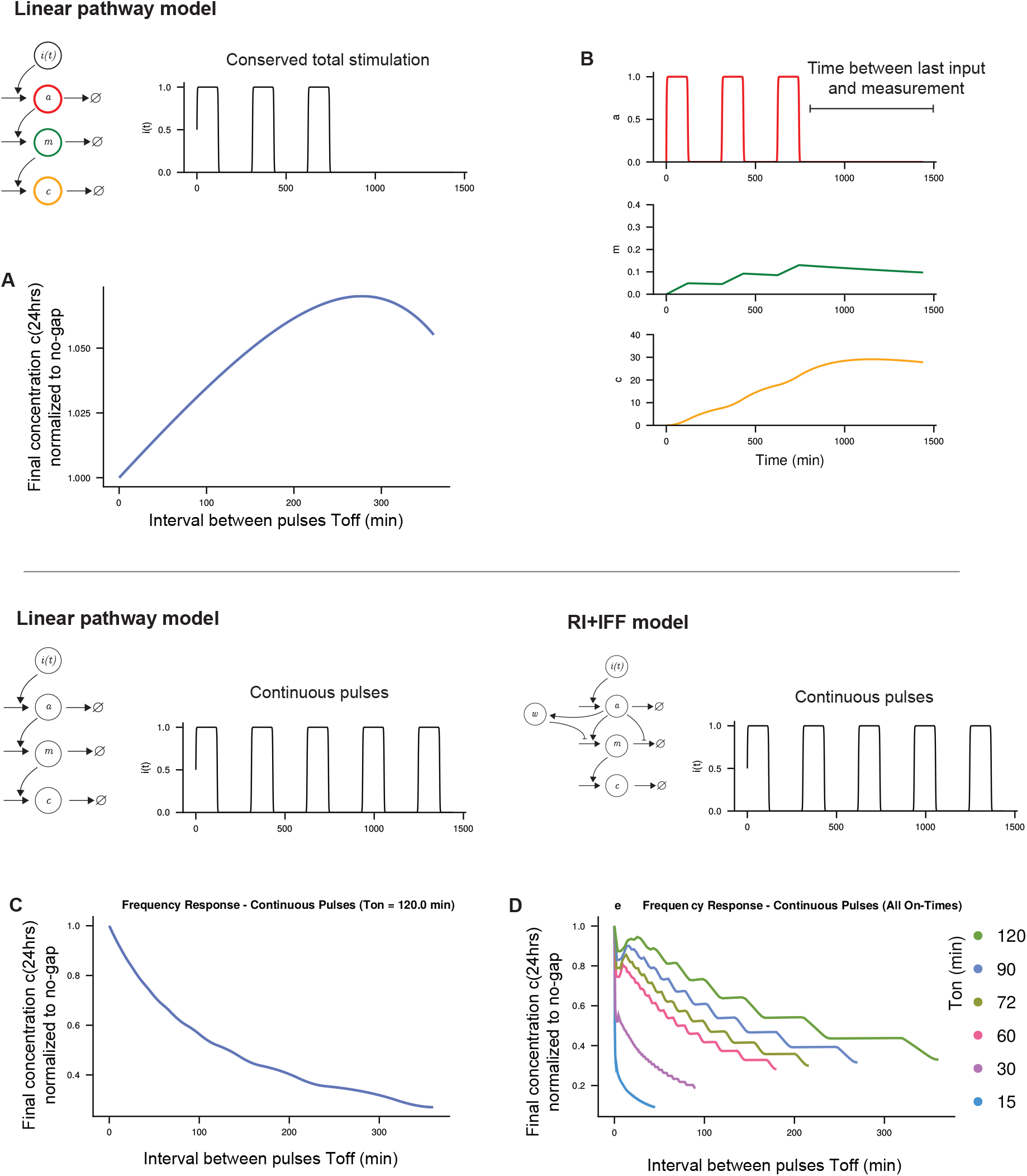
All models including the minimal model with a linear pathway can exhibit an increase in response when *T*_off_ *>* 0, compared to a single no-gap pulse. A Linear arrow architecture, with constant total stimulation (as in experimental data in (10)). B Given constant total stimulation time, longer gaps imply shorter time until the measurement, leading to increase in measured response. C Linear arrow architecture, but with continuous pulsatile input. D RI+IFF model with continuous pulsatile input, single on-time. E RI+IFF model with continuous pulsatile input, all on-times.

This raises the question of whether the real system would exhibit a true band-pass response, if pulses continue until measurement. While an experiment with continuing pulses is challenging, we can use the trained cross-validated RI+IFF model to simulate the T Cell’s response to continuing pulses. Note that the exploration of hypothetical continuous pulses in this section does not invalidate the results of previous sections: linear pathway model still fails to explain the experimentally-observed data.

### Validation on independent experimental data with distinct stimulus protocol

We next sought to test how generalizable our finding of the RI+IFF pathway is. An independent experiment by a different research group (11) used a LOVTRAP-based OptoCAR setup to present pulses of on-off T Cell Receptor binding to T cells, providing a potential independent validation opportunity for us. The protocol is different, for example, the researchers in (11) use continuous pulses and measure CD69 in a different manner (10, 11), and explored input pulses at fixed duty ratio *T*_on_*/*(*T*_on_ + *T*_off_) and period *T*_on_ + *T*_off_, as shown in Figure 6. We adjusted our model simulation and learning protocols, and retrained the RI+IFF model against this independent dataset. This involved changing the stimulus *i*(*t*) and modifying the final time to 22 hours (11).

**Figure 6.**
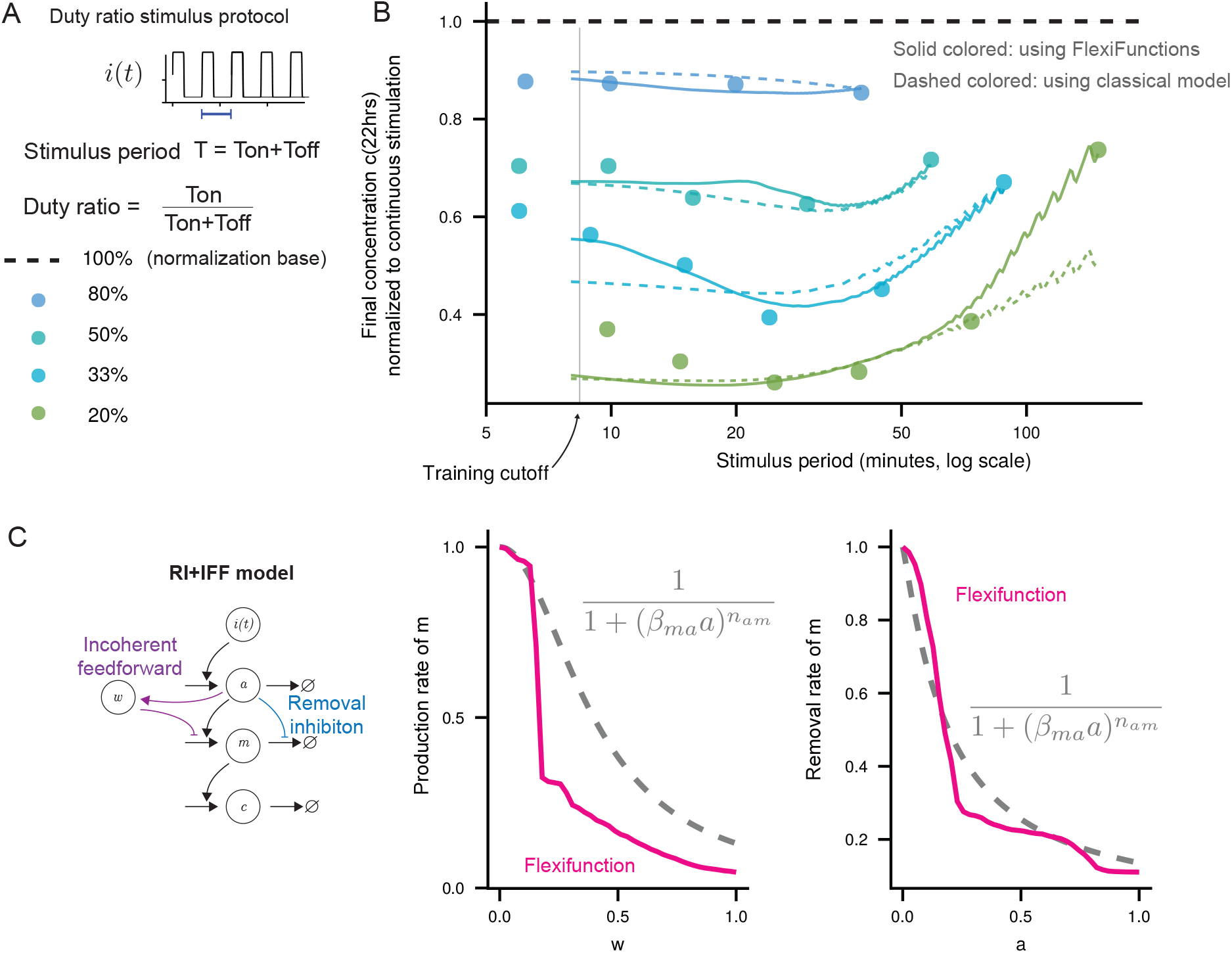
Validating the RI+IFF model on independent experimental dataset from (11) A Experimental protocol from (11) has continuous pulses, and parameterizes the stimulus protocols by stimulus period and duty ratio. B Band-stop behavior (a valley in cellular response at intermediate stimulus period) is exhibited by the experimental data (circles) and can be fit by the RI+IFF model (solid lines). The sharp zip-sags at long period in the model prediction result from measuring an oscillating signal at a fixed timepoint. The RI+IFF model with classical mass-action functions is shown as colored dashed lines. C Best-fit regulatory functions.

We found that the model fits the data to within root-mean-square error 0.021 (below the standard error of the mean in (10), our threshold for success), for stimulus periods above 8 minutes (lines in Figure 6). As expected, best-fit parameter values are slightly different, reflecting differences in experimental setups: *τ*_*a*_ = 8.3, *τ*_*m*_ = 3.0, *τ*_*w*_ = 106.4, *τ*_*c*_ = 10^6^ minutes. These are all the same order of magnitude as the best-fit parameter for the data in (10). Interestingly, the regulatory function for production rate of *m*, in incoherent feed-forward pathway, has a similar character as the regulators trained on the data from (10), as shown in Figure 6C, with sensitive dependence at low *w* becoming less sensitive at high *w*. Higher frequencie ranges were also explored in (11) (shorter stimulus period) compared to (10), as small a period as 2 minutes, smaller than the minimal on-time period of 15 minutes explored in the experiments of (10). The model was unable to fit the high-frequency data of (11), so we restricted to stimulus periodicity greater than 8 minutes. The inability to fit the high-frequency data could reflect additional nonlinearities or components acting at very short timescales — perhaps in the LOVTRAP experimental setup rather than the T Cell itself, as discussed above.

Despite differences in cell type, stimulation protocol, pulse durations, and measurement readout, the RI+IFF model is able to fit the data of (11) quantitatively. This striking result provides further confidence in identifying this pathway as a potential explanation for the temporal response of T Cell activation. Furthermore, the RI+IFF model with mass-action (a.k.a., classical, Hill-type) assumptions failed to fit the frequency response (dashed lines in Figure 6), as it failed for the Harris data above. This further supports our secondary result, that simple models may at first appear incapable of explaining complex data, but might indeed be able to, by relaxing functional form assumptions.

Note the small zig-zags in the frequency response at long periods (low frequency) at the right of Figure 6B. These are due to the measuring protocol, which is taken at a specific point in time, chosen in our simulation to match the protocol of (11). For an oscillating signal, this means that small changes in the oscillation period will result in an oscillating value of *c*(*t*) at a fixed time *t*.

### Model predicts non-saturated CD69 levels

One of the observations of the best-fit model time series in Figure 4 is that the output variable *c*(*t*), which we identify with CD69 surface expression level, does not saturate, meaning that it does not stop increasing if its upstream signal is present. Yet, biophysically, at some point, CD69 surface expression must saturate, so it is a meaningful model prediction that it has not saturated over the stimulus protocol and timeframe of the experiments of 24 hours. To confirm that this is a structural prediction of the model, rather than of a particular parameter regime, we repeated model learning with constrains on the model parameter *τ*_*c*_. This parameter, which has units of minutes, appears in Equation 18 and controls the timescale over which *c*(*t*) saturates. (Note that a small value of *τ*_*c*_ is a necessary condition for early saturation, but not a sufficient condition, since a slow timescale might be present in other upstream components.) We repeat model learning where *τ*_*c*_ is restricted to ten-fold intervals, and show the best-fit root-mean-square error in Figure 7. The model cannot be made to fit within experimental error bars for *τ*_*c*_ *<* 10^3^ minutes. So, a prediction of the model is that CD69 is not saturated under these experimental conditions. This prediction is perhaps unsurprising, since the range of CD69 readouts continues to change over the full set of protocols in both (10) and (11).

**Figure 7.**
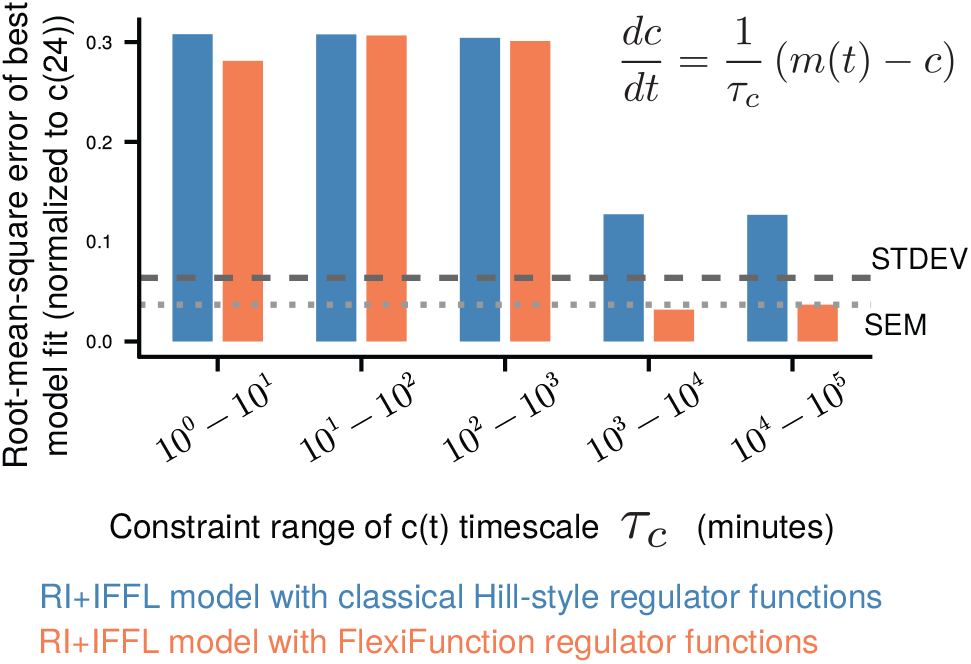
The timescale of *c*(*t*) must be large. The parameter *τ*_*c*_, in minutes, determines the characteristic timescale at which *c*(*t*) approaches its steady-state response to upstream pathway components, via Equation 18 (shown in the figure). Constraining the learning algorithm to ranges of values for *τ*_*c*_ results in worse root-mean-square errors.

## Discussion

The cross-validation we performed demonstrated that the model presented here — removal-inhibition and incoherent feed-forward, with nonlinear regulatory function — can predict datapoints withheld from training. This suggests it may be more generally predictive. If so, the model may have the ability to aid in the design of stimulus schedules for optimal T cell responses. For example, it could be used to design a stimulus schedule that maximizes activation, or minimizes activation, or maximizes it over a particular period before minimizing it. Similar predictive mathematical models have contributed to the design of schedules for administering cancer treatement drugs (52) and have been suggested for T-cell-based therapies (40).

Our findings yield several experimentally testable predictions. First, the model structurally requires that CD69 surface expression has not saturated over the 24-hour experimental window (Figure 7). Time-course measurements of CD69 during sustained stimulation should therefore show accumulation rather than a plateau. Second, the low-pass behavior observed in the data of (10) — in which pulsed stimulation with large gaps yields a stronger response than continuous stimulation — is predicted to be a consequence of the constant-total-stimulation-time protocol (Figure 5). If the experiment is repeated with pulses continuing until the measurement time, the model predicts that all pulsed conditions will produce a response at or below the no-gap baseline. Third, the best-fit refractory timescale *τ*_*w*_ ≈ 130 minutes predicts a specific recovery time from adaptation: after prolonged stimulation, gaps shorter than ~2 hours should produce little recovery of response strength, whereas gaps of ~6–7 hours (several multiples of *τ*_*w*_) should restore the full naive response amplitude.

A strength of phenotypic modeling is that the model components need not correspond to specific molecules. Nonetheless, we originally envisioned that the intermediate component *m* is a quantity related to CD69 mRNA or a transcriptional regulator thereof. If this identification is established, the model makes further testable predictions. In the time series of the best-fit model (Figure 4B), the first pulse of a rapid pulse train produces a markedly larger peak in *m* than subsequent pulses. A measurement of the corresponding molecular species should therefore show a distinctly elevated first-pulse response that subsequent pulses fail to match. Furthermore, because *τ*_*m*_ ≈ 1–3 minutes while *τ*_*c*_ → ∞, this intermediate quantity should decay within minutes of stimulus removal, even as CD69 surface protein persists for hours.

The phenotypic model revealed by pulsatile stimulus contains two common motifs: removal-inhibition of an intermediate component, and an incoherent feed-forward loop. Removal-inhibition provides the ability to respond rapidly to changes (26). This motif gives rise to band-stop behavior, which has been suggested to allow T cells to filter out interactions with healthy cells, to allow more effort to focus on its targets (11). Incoherent feed-forward motifs are found throughout biology (26, 39) and endow systems with adaptation: they become desensitized following prolonged exposure, effectively redefining their baseline.

When a model fails to fit every feature of a dataset, including small quantitative features that do not change the categorical behavior, it is tempting to add model complexity. The modeling approach presented here allows the same, interpretable arrow diagram model to be given additional quantitative flexibility to be learned from data. The method is similar in spirit to a class of approaches in which the right-hand-side of a dynamical model is replaced with a universal function (34–36), with a few key distinctions. First is our viewpoint that monotonic-increasing or monotonic-decreasing connections between nodes are desirable for interpretability. Second is that all the universal functions we used were single-input, single-output. Indeed, the approach can adapt to the specific project’s definition of interpretability. The approach could be repeated with non-monotone regulatory functions — if your objective of interpretability were simply the connections between components. Or, it could be repeated with more constrained functions, for example functions that have log-derivatives no larger than 1.0, ensuring no arrow exhibits ultrasensitivity, and any ultrasensitive response arises from the arrow diagram itself, not individual arrows. This is connected to another distinction between our approach and similar approaches, which is our use of piece-wise linear approximation to represent the universal function. Imposing the monotonicity constraint proved most computationally efficient for the piecewise-linear universal function approximator (Equation 21) compared to Chebyshev expansions, Fourier series, Taylor series, and multi-layer perceptrons. Indeed, early versions of our code used multilayer perceptrons but the piece-wise linear version provides a computationally cheaper and, in our view, more elegant implementation for this specific application.

In this work, we manually explored the space of possible arrow-diagrams, represented visually as moving left-to-right in the table in Figure 3A. For models with the complexity explored here, with 4 internal dynamic components, each pair linked bidirectionally by either promotion, inhibition, or neither, the set of all possible arrow sets with 4 components is 3^(4 choose 2) = 729. For models that are cheaper to evaluate, there are existing methods for exhaustive, systematic search of model-space (4, 44, 53–55), but these are presently suited to models with more restricted functional form assumptions that those that proved necessary here. For this dataset, a surprisingly simple model proved able to fit the data with the desired root-mean-square residual, making manual ad-hoc model exploration feasible. However, future data sets may necessitate combining the flexifunction approach with automated exploration of the space of arrow-diagrams.

## Methods

All code to define, simulate, and learn models is provided at: https://github.com/allardjun/PulsatileModelLearning

### Models

The mathematical models were scaled (non-dimensionalized) in quantity value, while time was kept in physical units of minutes. In all cases, *i*(*t*) represents the input signal of optically-activated TCR, either 0 or 1 at any time *t*. We identify *c*(*t*) as the surface expression of CD69, in nondimensional units so that only comparisons of *c*(*t*) are meaningful. Model equations were constrained so that if *i*(*t*) = 0 then the steady-state *c*(*t*) = 0 in agreement with (10). When rescaling, there are two mathematically equal choices of whether the *τ*_*i*_ multiplies the full right-hand side or just the linear decay term. We found that when fitting data, *τ*_*c*_ was larger than the other *τ*_*i*_’s, so for numerical integration stability and accuracy, we rescaled *c* with *τ*_*c*_ as shown below.

### Linear architecture

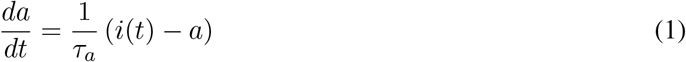

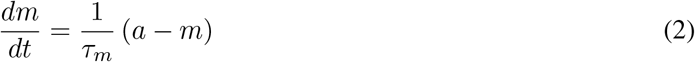

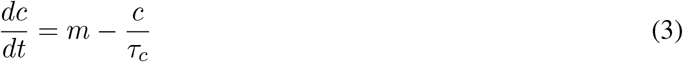

Parameters: *τ*_*a*_, *τ*_*m*_, *τ*_*c*_.

### Linear architecture with explicit LOVTRAP kinetics

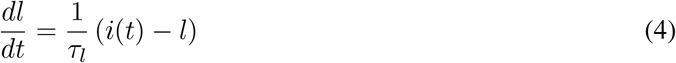

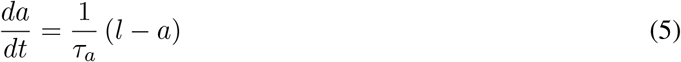

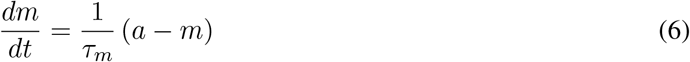

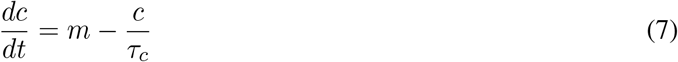

Parameters: *τ*_*l*_, *τ*_*a*_, *τ*_*m*_, *τ*_*c*_.

### Removal-Inhibition (RI)

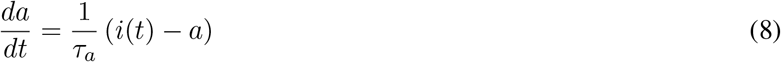

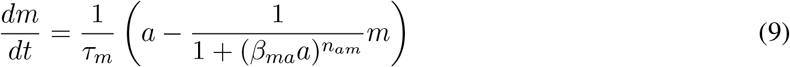

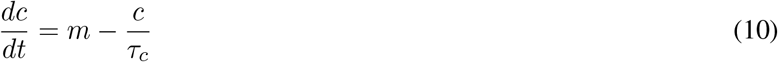

Parameters: *τ*_*a*_, *τ*_*m*_, *τ*_*c*_, *β*_*ma*_, *n*_*am*_.

### Incoherent Feed-forward (IFF)

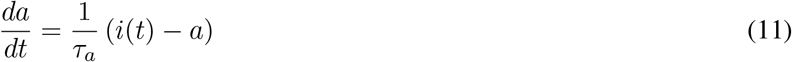

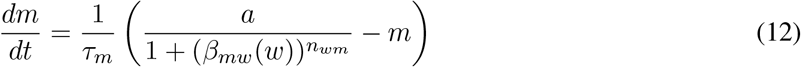

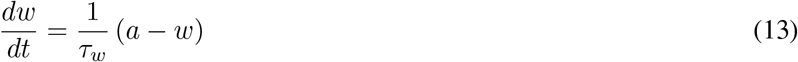

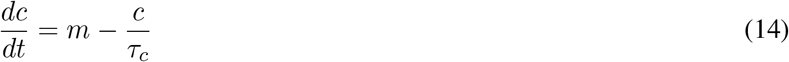

Parameters: *τ*_*a*_, *τ*_*m*_, *τ*_*c*_, *τ*_*w*_, *β*_*mw*_, *n*_*wm*_.

### Removal-Inhibition and Incoherent Feed-forward Loop (RI+IFF)

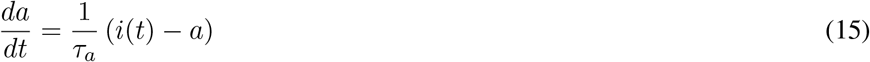

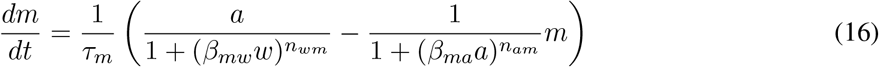

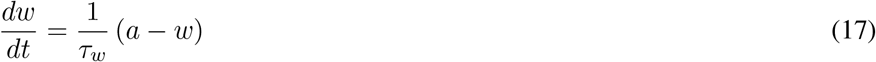

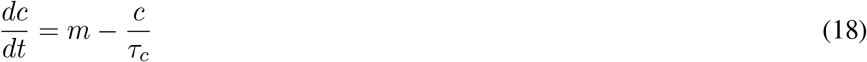

Parameters: *τ*_*a*_, *τ*_*m*_, *τ*_*c*_, *τ*_*w*_, *β*_*mw*_, *n*_*wm*_, *β*_*ma*_, *n*_*am*_.

### Generation of frequency response plot for comparison with data

The experimental data from (10) consists of CD69 protein expression measurements in engineered T cells subjected to pulsatile optical stimulation with varying pulse durations (*T*_on_ = 15, 30, 60, 72, 90, 120 minutes) and interpulse intervals (*T*_off_ = 0–480 minutes).

To match these conditions, we implement a pulsatile input function that generates time-dependent stimulation patterns with constant time-integral across different pulsing protocols. This constraint ensures that different pulse/interval combinations deliver equivalent total stimulation, matching the experimental design where pulse amplitude was adjusted to maintain constant integrated dose. Each experimental condition (6 × 14 = 84 total conditions) is simulated by integrating the ODE system with the corresponding pulsatile input, producing model predictions for the 24-hour CD69 readout that can be directly compared to experimental measurements.

Frequency response analysis provides quantitative comparison between model predictions and experimental data across the full parameter space of pulsing conditions. For each model and parameter set, we systematically simulate all 84 experimental conditions by solving the ODE system with appropriate pulsatile inputs using high-precision numerical integration (Vern9 algorithm with adaptive tolerance).

The final output concentration *c*(24h) serves as the model prediction for CD69 expression at the experimental endpoint. To match experimental normalization, all model outputs are normalized to the first off-time condition within each on-time series:

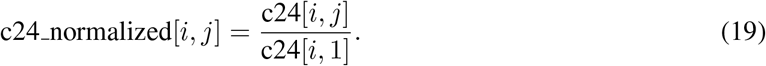

This produces a 6 × 14 frequency response matrix that directly corresponds to the experimental data structure. Model-data comparison is visualized through heatmaps and line plots showing normalized CD69 expression as a function of interpulse interval for each pulse duration. The root-mean-square (rms) error between model and data serves as the primary metric for model performance:

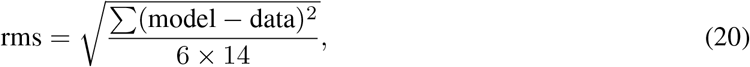

providing normalized error estimates across all experimental conditions.

### Flexifunctions implementation

Flexifunctions provide a data-driven approach to extend classical mechanistic models by replacing classical functions with learnable functions. Each flexifunction is parameterized by a *n*_dofs_ parameters, *θ*_1_..*θ*_*n*dofs_, that define breakpoints for monotonic interpolation over the unit interval [0, 1].

The function is built using cumulative sums of squared parameters,

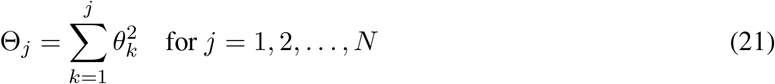

and Θ_0_ = 0. This guarantees the Θ_*j*_ values are increasing and have zero-intercept. The piecewise linear interpolation occurs between evenly-spaced adjacent cumulative values,

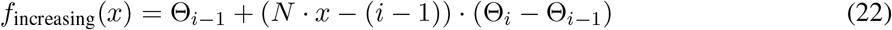

where *i* = min(⌊*N* · *x*⌋ + 1, *N*) is the interval index. Note the optimization algorithm can freely adjust *θ*_*k*_ ∈ ℝ without violating the constraints (increasing and zero-intercept). To obtain a continuous, strictly-positive, decreasing function with intercept at unity, we take

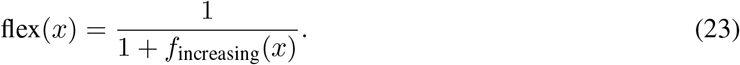

Thus, these functions satisfy

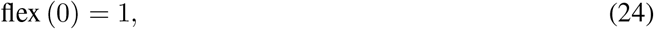

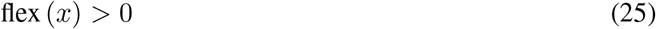

And

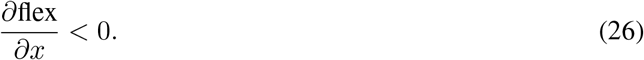

As *n*_dofs_ → ∞, the piece-wise linear functions captured in this basis approach the set of arbitrary decreasing functions subject to Equations 24-26. These monotonic-increasing, zero-intercept functions are then inserted into the right-hand side of differential equations, for example for the RI+IFF model,

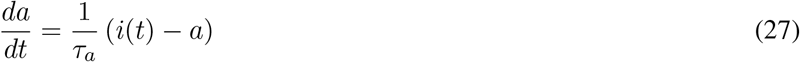

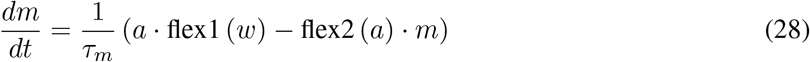

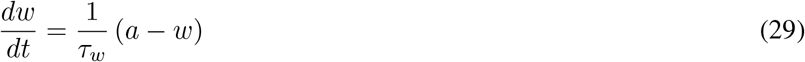

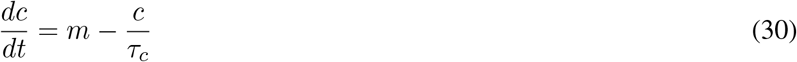

Unless otherwise noted, flex1 and flex2 each contain *n*_dofs_ parameters, where *n*_dofs_ = 64 as determined by cross-validation, see below.

### Learning Algorithm

The framework computes rms errors compared to experimental between-replicate variability (standard deviation for 3 replicates of 0.0638, so standard error of the mean is 0.0368). The codebase uses a dual parameter representation system to ensure all optimization parameters are well-behaved across the entire real number line (™ ∞, +∞). Before parameters are visible to the optimization algorithms, they are transformed into optimization-friendly “represented” parameters using log and square-root transformations. After optimization, they are “derepresented” back to physically meaningful values that appear in the ODEs above.

### Learning Algorithm for Classical Models

Classical models are optimized using global optimization methods. The primary algorithm is the adaptive differential evolution variant `BBO adaptive de rand 1 bin()` from the `Optimization.jl` Julia package, which performs global parameter space exploration for the 3–8 classical parameters.

The algorithm converges by 2.5 × 10^4^ iterations (Supplemental Figure S6).

### Learning Algorithm for Models Containing flexifunctions

Flexi models present a significantly more challenging optimization landscape due to the high-dimensional parameter space (20–256 flexifunction parameters plus classical parameters). The learning protocol implements a three-stage “corduroy” optimization strategy that alternates between different parameter subsets to manage complexity and avoid local optima. An example learning trajectory is shown in Supplemental Figure S6.

First, we run the corresponding classical model as above to generate an initial guess. The corduroy learning protocol then alternates between CMA-ES optimization of flexifunction parameters (holding classical parameters fixed) and a simplex refinement (Rowan’s Subplex algorithm (56)) of classical parameters (holding flexifunctions fixed). CMA-ES (Covariance Matrix Adaptation Evolution Strategy) is particularly well-suited for flexifunction optimization due to its adaptive covariance matrix that captures parameter correlations and its robust performance in high-dimensional spaces. Interestingly, the simplex iterations decrease the loss only slightly (not visible in Supplemental Figure S6), and yet are sufficient to allow the following CMA-ES iterations to discover a significant improvement, even though the previous CMA-ES round of iterations had converged.

The algorithm maintains adaptive step-size control (*σ*_0_) computed based on parameter bounds and implements custom hyperparameter schedules for population size and termination criteria. Multiple corduroy rounds ensure convergence, with each iteration building upon improved parameter estimates from the previous round. This approach effectively decomposes the complex joint optimization into manageable subproblems while maintaining global search capabilities for the high-dimensional flexifunction parameter space.

We find that 120 CMA-ES iterations, 200 simplex iterations, each repeated 5 times leads to unchanging solutions and appears converged (Supplemental Figure S6). This procedure takes around 72 hours on Intel Xeon Gold 6230 with 8GB of memory.

Note that the optimization function is differentiable, but not convex, so gradient descent methods fail to improve the fit.

### Cross-validation and degrees-of-freedom selection

Flexifunction complexity is determined through systematic cross-validation to balance model expressiveness against overfitting. We implement *k*-fold cross-validation with *k* = 7 (since the number of data points is divisible by 7). This partitions the 84 experimental conditions into randomly-selected but non-overlapping training and validation sets, across all pulsing conditions. For each *n*_dofs_ value (2, 4, 8, 16, 32, 64, 128, 256), for each of the 7 random subsets, we train flexi models on 72 conditions and evaluate performance on the remaining 12 conditions, rotating through all fold combinations. The validation metric is RMS error computed on held-out data. This process generates learning curves showing training and validation error as functions of model complexity, with the optimal *n*_dofs_ selected as the point where validation error is minimized before overfitting occurs.

As shown in Supplemental Figure S4, the fit quality generally improved as more degrees of freedom were added to the flexifunctions. Then, the root-mean-square error was measured on the withheld subset. The error in the testing subset exhibits a pronounced minimum around *n*_dofs_ = 32 degrees of freedom. Note importantly that adding more degrees of freedom did not allow unlimited improvements to the fit even on the training set, but rather plateaus at large *n*_dofs_. This suggests that these models have structural constraints that would allow a model to be rejected, even with infinite degrees of freedom.

### Testing the learning algorithms and flexifunction approach on synthetic data

To study the performance of the learning algorithm, and ensure correctness of the implementation, we apply the setup to synthetic data in which the ground truth is known. We simulate two ground truth models: The first is RI+IFF model with classical Hill function for the two regulatory functions, as described in Equations 15-18. The second is the RI+IFF, so the model described in Equations 27-30, where the regulatory function are non-Hill, non-sigmoidal “wobbly” functions. To achieve this, we selected functions that are monotonic decreasing, satisfy the correct left boundary constraint, but have sufficient non-Hill character. The functions we chose are

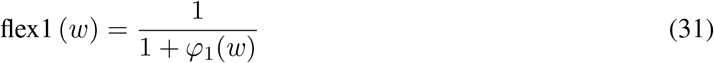

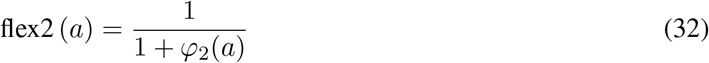

and

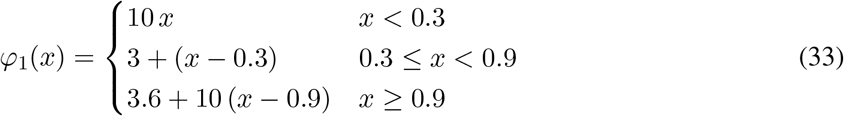

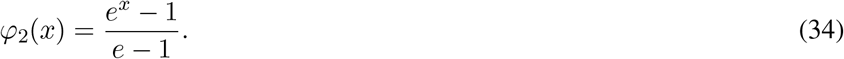

These models are the most complicated models we study in this work, and the distinction between them the most subtle, making this the ideal test of our implementation. For each of those two ground truth models, we generate synthetic data at 3 noise levels: 10% of the Harris standard error of the mean (SEM), 1x the Harris SEM, and 2x the Harris SEM. As a success criteria, we use the same as used in the real data analysis: the model must fit to within the mean SEM of the data (so, 10%, 100% and 200% of the Harris data). With this synthetic data as the training target, we perform learning using two models: the RI+IFF model with classical Hill functions, and the RI+IFF model with flexifunctions for the two regulatory functions.

Results are shown in Figure S7. For classical ground truth, both models fit within the noise. For nonlinear regulatory function model, at the experimental standard error of the mean, the classical model has rms error larger than the noise, while the fleximodel has rms error below the noise. Thus, the model learning pipeline is able to distinguish models at the experimentally-observed noise level.

## Acknowledgements

We thank Omer Dushek, Brady Berg, Katie Bogue, Jack Corrette, Austin Marcus for useful discussion. This work was supported by NSF DMS 2052668.

## 1 Supplemental material

**Figure S1:**
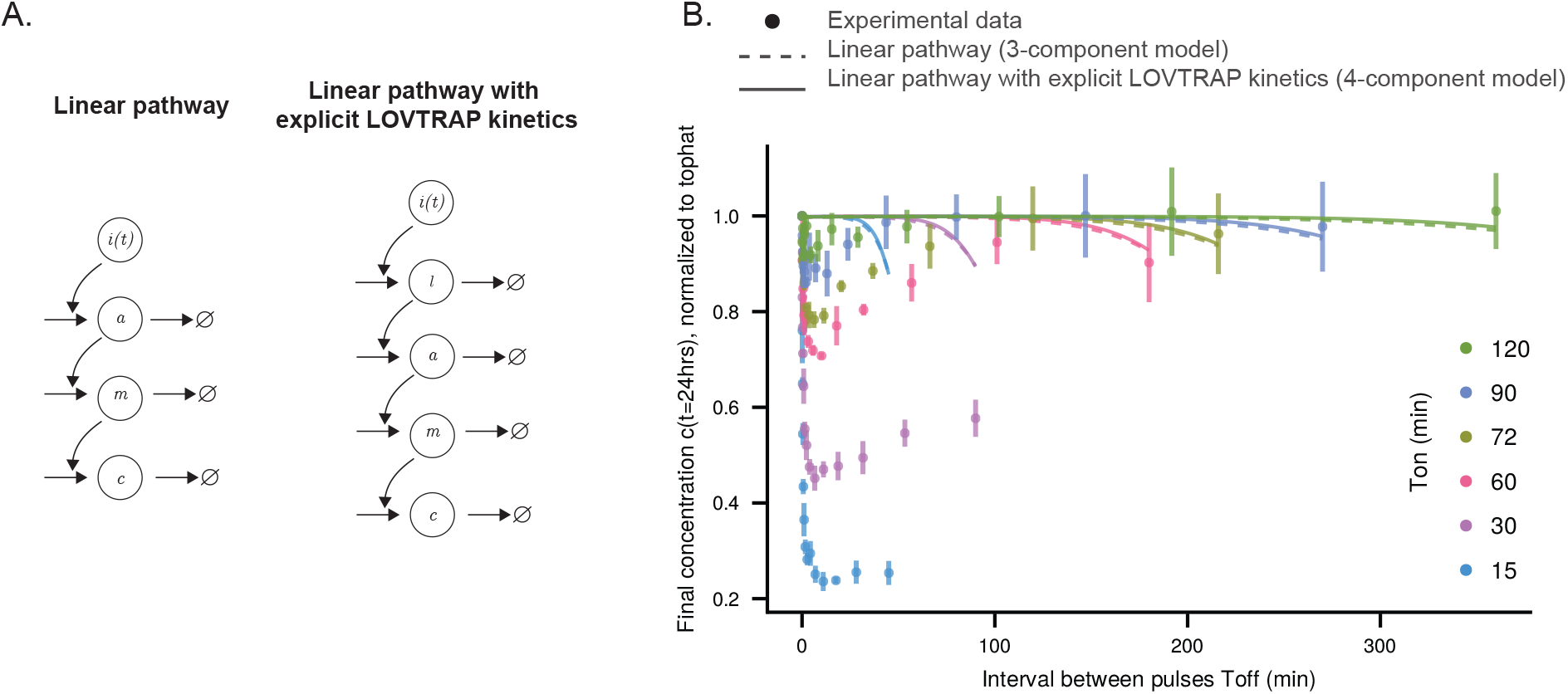
Linear pathway models, with or without explicit LOVTRAP dynamics, fail to explain the frequency response data. A Pathway schematics for linear pathway and linear pathway with explicit LOVTRAP kinetics. B Best-fit frequency response of linear pathway models. Solid: Linear pathway with explicit LOVTRAP dynamics (4-component model); dashed: linear pathway (3-component model). Plot symbols show data from (10).

**Figure S2:**
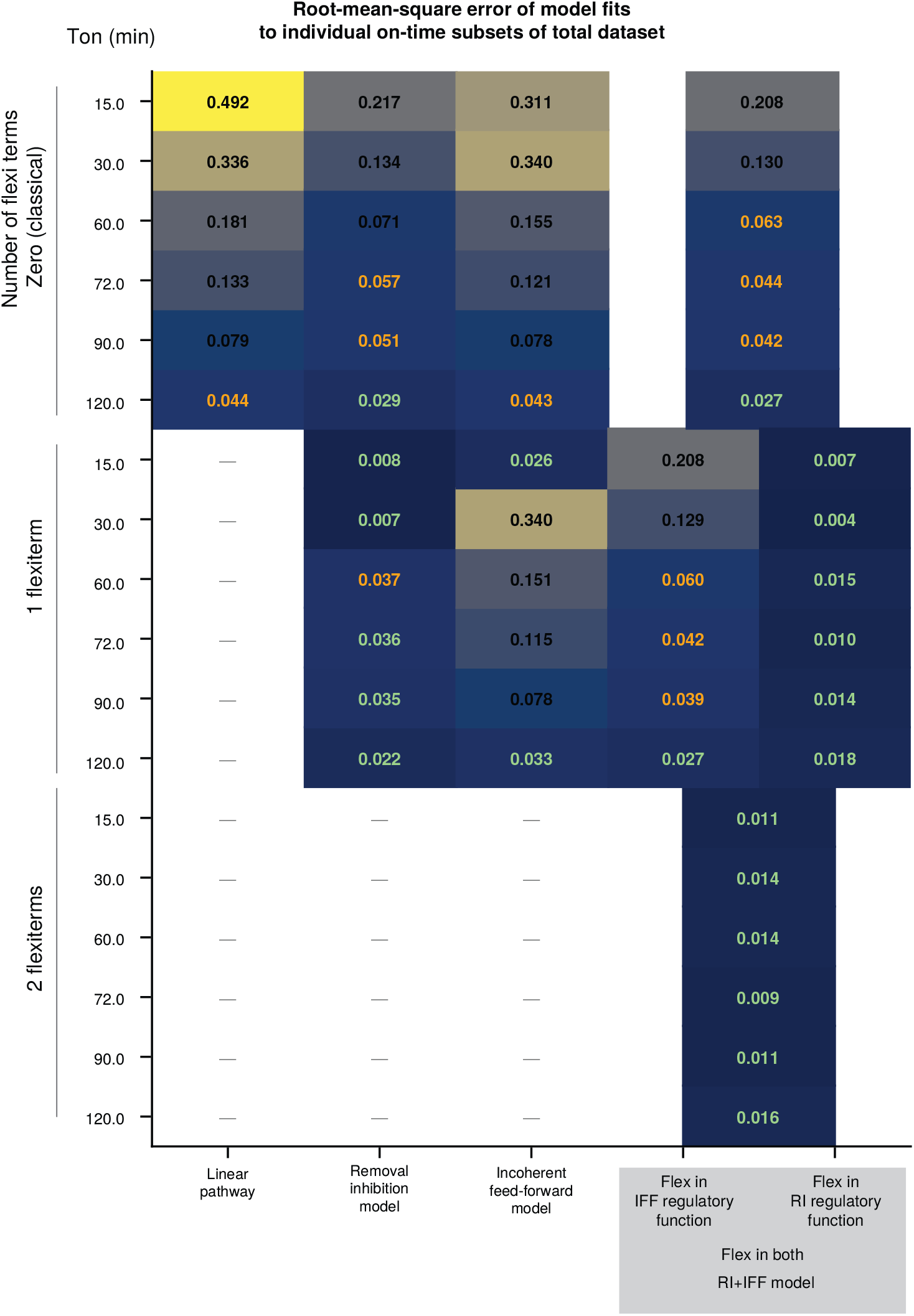
Best-fit rms error of models, fit to individual on-time series, for models with classical mathematical biology functional forms, and 1 or 2 flexifunctions. Text font indicates whether the fit root-mean-square error was smaller than the experimental data’s standard deviation (orange) or smaller than the experimental data’s standard error of the mean (green). Note that the RI+IFF model, which has both removal-inhibition and incoherent-feedforward regulatory functions, there are two options for where to put a single flexifunction.

**Figure S3:**
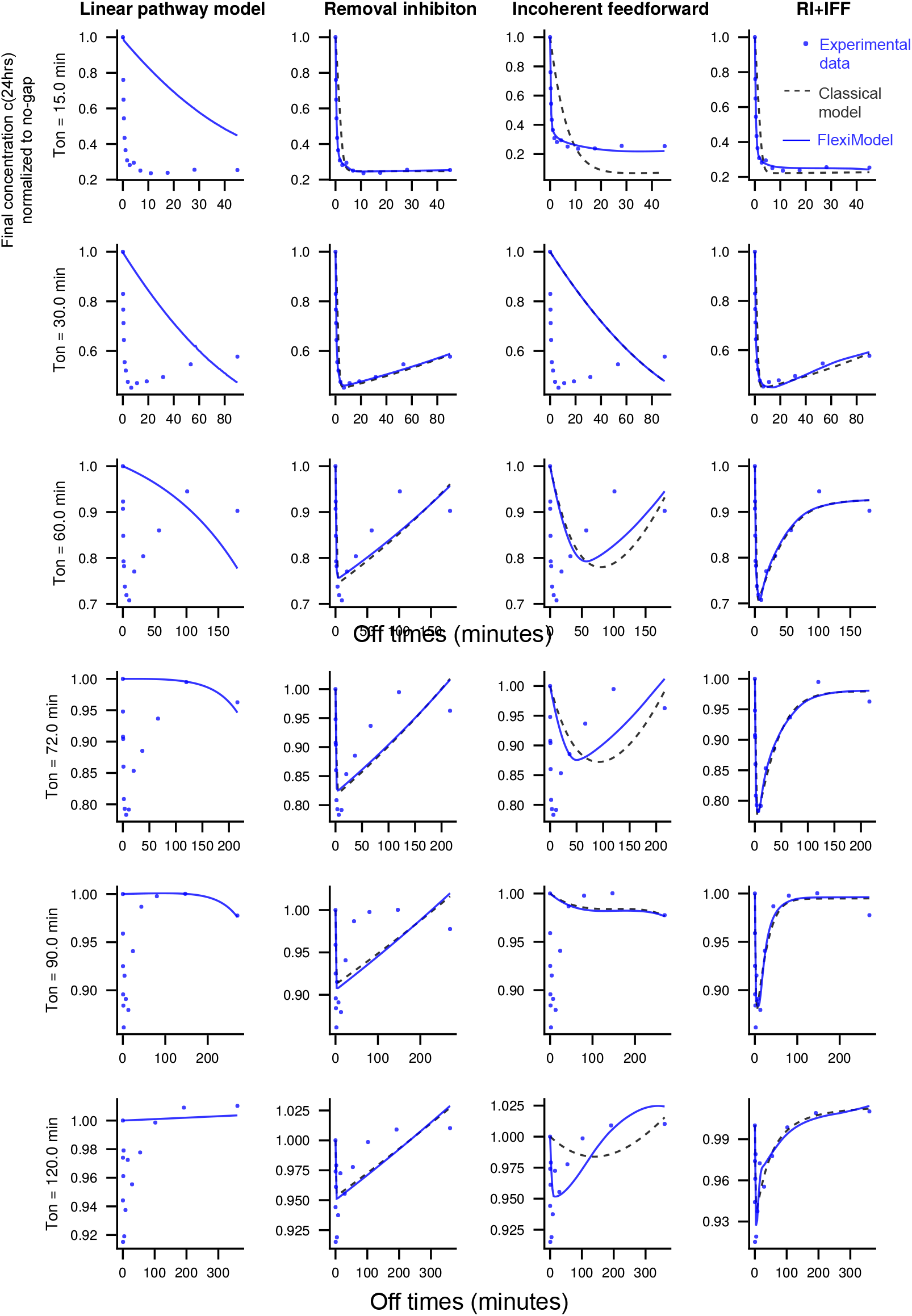
Best-fit rms error of models, fit to individual on-time series, for models with classical mathematical biology functional forms (for 4 models), and 1 flexifunction (for 3 models). These correspond to a subset of rms errors reported in Supplemental Figure S2.

**Figure S4:**
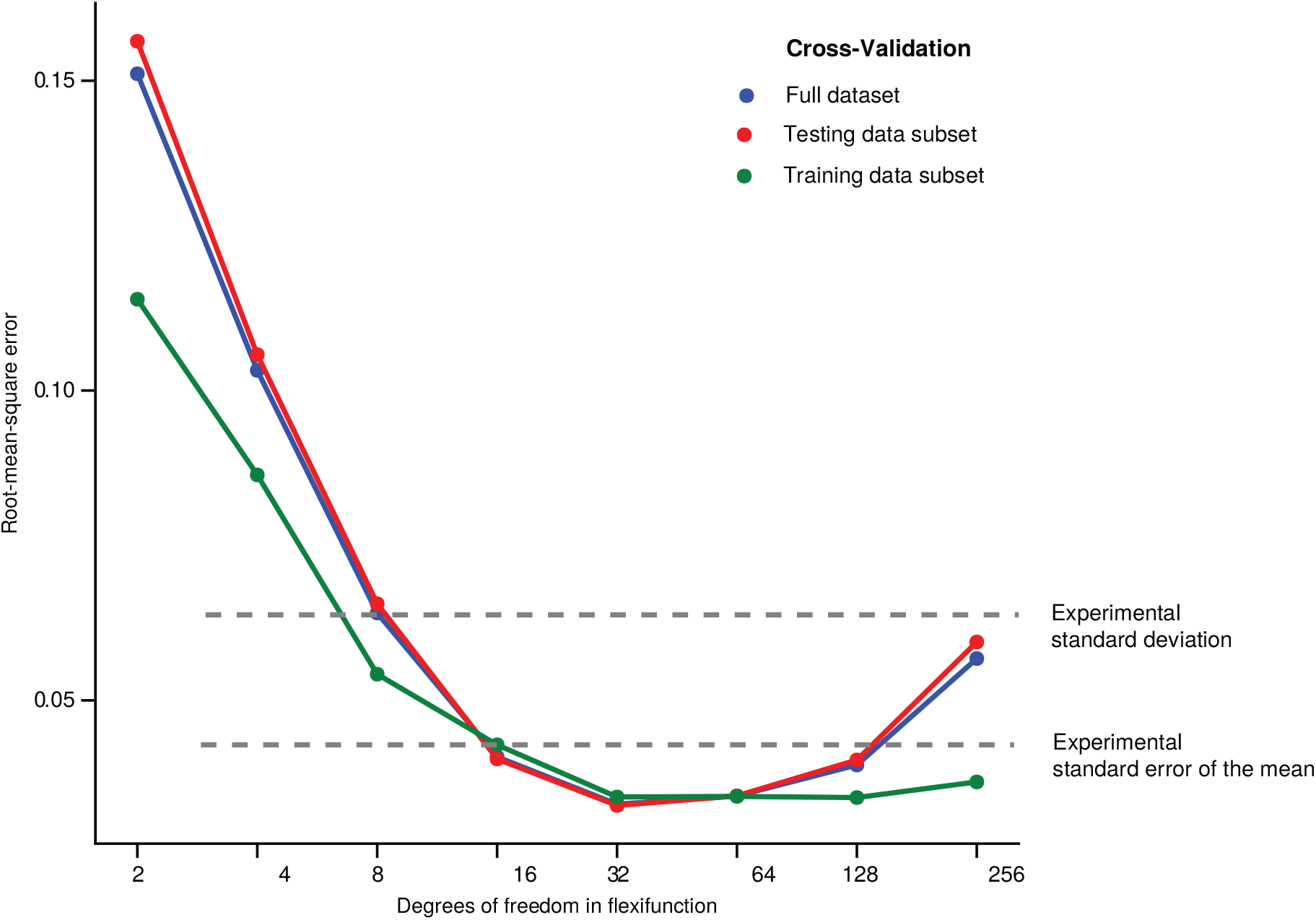
Cross-validation analysis allows choice of degrees of freedom in flexifunctions to avoid over-fitting. RI+IFF model (two flexifunctions) was fit to subsets of the full dataset, specifically 7 subsets were generated where 14 datapoints were removed without replacement (7 was chosen since the total number of datapoints was 7 × 14). The fit quality, measured by root-mean-square error to the training set (green), generally improved as more degrees of freedom were added to the flexifunctions. Then, the root-mean-square error was measured on the removed subset (testing data subset, red). The error in the testing subset exhibits a pronounced minimum around 32 degrees of freedom. Note importantly that adding more degrees of freedom did not allow unlimited improvements to the fit to the training set — this suggests that these models have structural constraints that would allow a model to be rejected, even with infinite degrees of freedom. The error on the training data is slightly above the testing data for 2 points, which may be due to stochastic optimization. The slight increase in error on the training data at very large degrees of freedom may indicate the algorithm was not yet converged at these very large model sizes.

**Figure S5:**
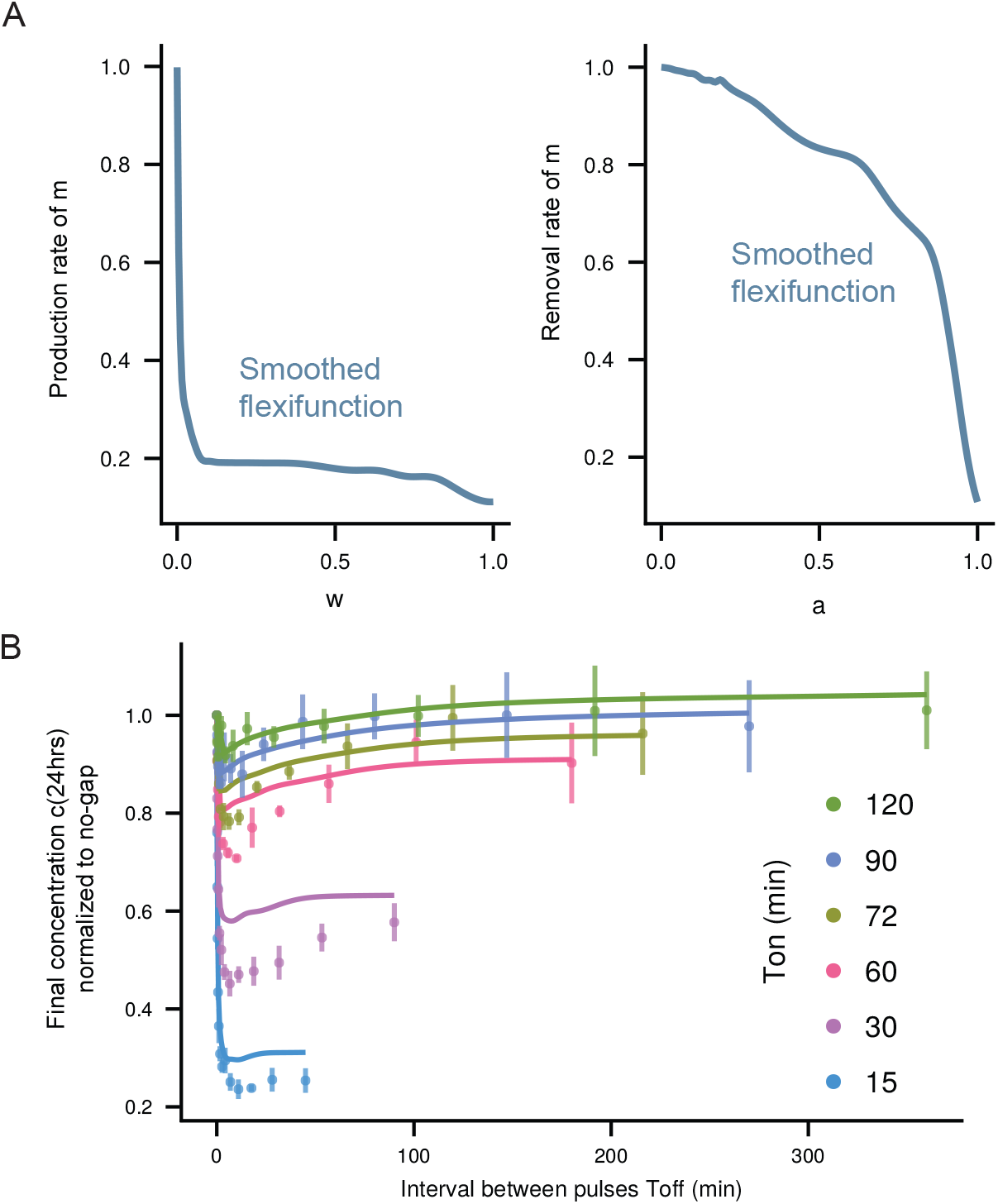
The regulator functions learned from data can be smoothed, leading to minimal impact on qualitative and quantitative consistency with data. A Smoothed regulator functions. Compare with Figure 3B. B The impact of smoothed regulator functions on the predicted frequency response. Note the model was not retrained, so this is a true test of model sensitivity to details of the regulator functions.

**Figure S6:**
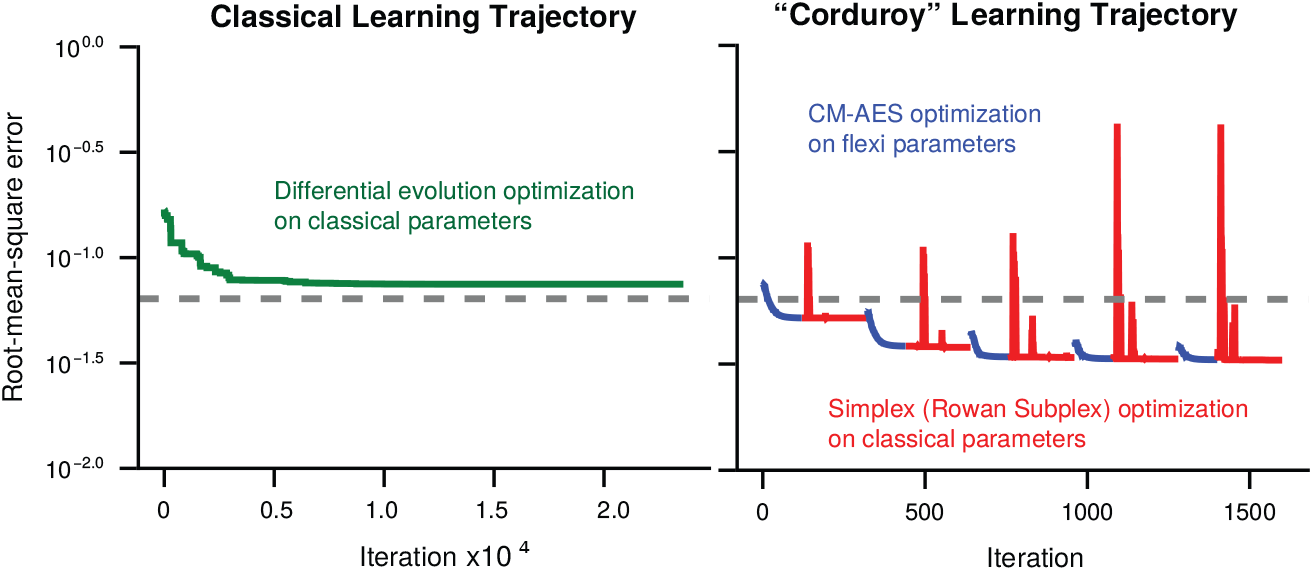
Iterative learning algorithm. To find the optimal parameter set, we first perform differential evolution optimization on just the classical model (left). Then, use these as initial guesses and repeat CMA-ES optimization on only the flexifunction parameters, keeping the classical parameters fixed, and simplex (Rowan’s Subplex) on just the classical parameters, keeping the flexifunction parameters fixed (right)

**Figure S7:**
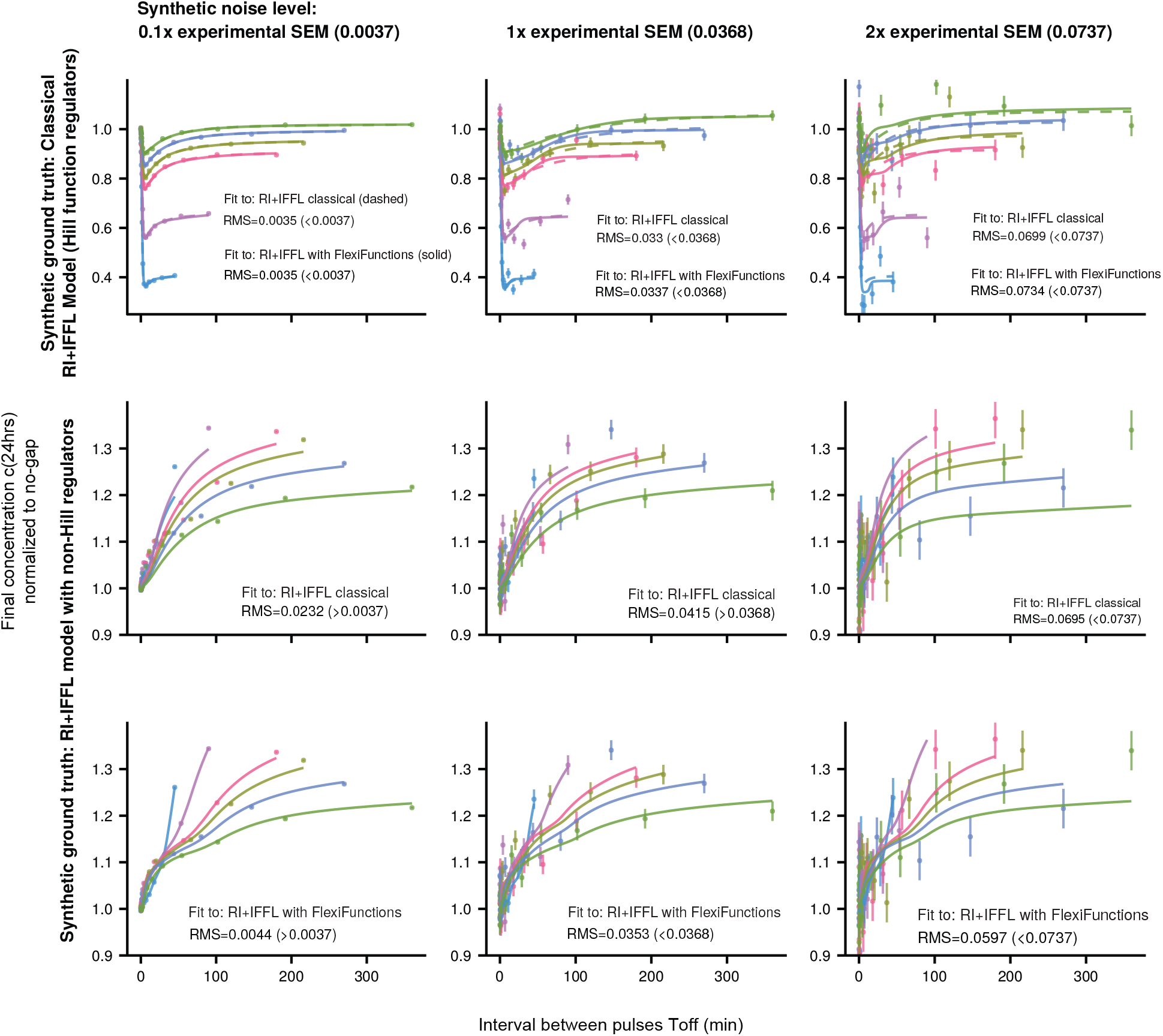
Results of learning on synthetic data in which the ground truth is known. The ground truth is classical RI+IFF model with Hill functions, Equations 15-18 (top row) or RI+IFF model with a non-Hill functions Equation 33-34 for the regulator (middle and bottom row), with three noise levels (columns), corresponding to 10%, 100% and 200% the standard error of the mean experimentally reported in (10). In each of those 6 synthetic datasets, the classical RI+IFF model with Hill functions is fit (top and middle rows), and the RI+IFF model with flexifunction regulators is fit (top and bottom rows). For classical ground truth, both models fit within the noise. For nonlinear regulatory function model, at the experimental standard error of the mean, the classical model has RMS error larger than the noise, while the fleximodel has RMS error below the noise. The family of curves represent different *T*_on_, with the same color legend as Figure 3

## Notes

### Competing Interest Statement

The authors have declared no competing interest.

### Summary of Updates

Two new results sections, one validating the model on an independent dataset, and the other demonstrating the model requires non-saturating dynamics in the output variable. Edits to test to improve writing and background.

## References

1. Kramer, J., Merwe, P.A. v. d. & Dushek, O. Platforms for studying cell-cell recognition by immune cells. Immunology and Cell Biology 103, 636–647 (2025).

2. Lever, M., Maini, P. K., Merwe, P.A. v. d. & Dushek, O. Phenotypic models of T cell activation. Nature Reviews Immunology 14, 619–629 (2014).

3. Makaryan, S. Z. & Finley, S. D. Enhancing network activation in natural killer cells: predictions from in silico modeling. Integrative Biology 12, 109–121 (2020).

4. Lever, M. et al. Architecture of a minimal signaling pathway explains the T-cell response to a 1 millionfold variation in antigen affinity and dose. Proceedings of the National Academy of Sciences 113, E6630–E6638 (2016).

5. Wertheim, K. Y. et al. A multi-approach and multi-scale platform to model CD4+ T cells responding to infections. PLoS Computational Biology 17, e1009209 (2021).

6. Kondo, T. et al. Engineering TCR-controlled fuzzy logic into CAR T cells enhances therapeutic specificity. Cell 188, 2372–2389.e35 (2025).

7. Patel, A. et al. Using CombiCells, a platform for titration and combinatorial display of cell surface ligands, to study T-cell antigen sensitivity modulation by accessory receptors. The EMBO Journal 43, 132–150 (2024).

8. Bechhoefer, J. Control Theory for Physicists (Cambridge University Press).

9. Szischik, C. L., Szemere, J. R., Balderrama, R., Vega, C.S. d. l. & Ventura, A. C. Transient frequency preference responses in cell signaling systems. npj Systems Biology and Applications 10, 86 (2024).

10. Harris, M. J., Fuyal, M. & James, J. R. Quantifying persistence in the T-cell signaling network using an optically controllable antigen receptor. Molecular Systems Biology 17, MSB202010091 (2021).

11. ODonoghue, G. P. et al. T cells selectively filter oscillatory signals on the minutes timescale. Proceedings of the National Academy of Sciences 118, e2019285118 (2021).

12. Schmid, E. M. et al. Size-dependent protein segregation at membrane interfaces. Nature Physics 12, 704–711 (2016).

13. Taylor, R., Allard, J. & Read, E. L. Simulation of receptor triggering by kinetic segregation shows role of oligomers and close contacts. Biophysical Journal 121, 1660–1674 (2022).

14. Zhang, Y. et al. The Influence of Molecular Reach and Diffusivity on the Efficacy of Membrane-Confined Reactions. Biophysical Journal 117, 1189–1201 (2019).

15. Clemens, L. et al. Determination of the molecular reach of the protein tyrosine phosphatase SHP-1. Biophysical Journal 120, 2054–2066 (2021).

16. Goyette, J. et al. Dephosphorylation accelerates the dissociation of ZAP70 from the T cell receptor. Proceedings of the National Academy of Sciences 119, e2116815119 (2022).

17. Liu, K. et al. Hydrodynamics of transient cell-cell contact: The role of membrane permeability and active protrusion length. PLoS Computational Biology 15, e1006352 (2019).

18. Clemens, L., Dushek, O. & Allard, J. Intrinsic Disorder in the T Cell Receptor Creates Cooperativity and Controls ZAP70 Binding. Biophysical Journal 120, 379–392 (2021).

19. Hausser, J., Mayo, A., Keren, L. & Alon, U. Central dogma rates and the trade-off between precision and economy in gene expression. Nature Communications 10, 68 (2019).

20. Nika, K. et al. Constitutively Active Lck Kinase in T Cells Drives Antigen Receptor Signal Transduction. Immunity 32, 766–777 (2010).

21. Dutta, D. et al. Recruitment of calcineurin to the TCR positively regulates T cell activation. Nature Immunology 18, 196–204 (2017).

22. Fernandez-Aguilar, L. M., Vico-Barranco, I., Arbulo-Echevarria, M. M. & Aguado, E. A Story of Kinases and Adaptors: The Role of Lck, ZAP-70 and LAT in Switch Panel Governing T-Cell Development and Activation. Biology 12, 1163 (2023).

23. Davidson, D., Bakinowski, M., Thomas, M. L., Horejsi, V. & Veillette, A. Phosphorylation-Dependent Regulation of T-Cell Activation by PAG/Cbp, a Lipid Raft-Associated Transmembrane Adaptor. Molecular and Cellular Biology 23, 2017–2028 (2003).

24. Manz, B. N. et al. Small molecule inhibition of Csk alters affinity recognition by T cells. eLife 4, e08088 (2015).

25. Bourassa, F. X. P., Achar, S., Altan-Bonnet, G. & Francois, P. Learning the principles of T cell antigen discernment. arXiv (2025). 2511.18626.

26. Alon, U. An Introduction to Systems Biology: Design Principles of Biological Circuits. Chapman & Hall/CRC Computational Biology Series (CRC Press,2020). URL https://books.google.com/books?id=udihDwAAQBAJ.

27. Kaplan, S., Bren, A., Dekel, E. & Alon, U. The incoherent feed-forward loop can generate nonmonotonic input functions for genes. Molecular Systems Biology 4, MSB200843 (2008).

28. Prybutok, A. N., Cain, J. Y., Leonard, J. N. & Bagheri, N. Fighting fire with fire: deploying complexity in computational modeling to effectively characterize complex biological systems. Current Opinion in Biotechnology 75, 102704 (2022).

29. Shen, C. et al. Differentiable modelling to unify machine learning and physical models for geosciences. Nature Reviews Earth & Environment 4, 552–567 (2023). 2301.04027.

30. Noordijk, B. et al. The rise of scientific machine learning: a perspective on combining mechanistic modelling with machine learning for systems biology. Frontiers in Systems Biology 4, 1407994 (2024).

31. Yu, R. & Wang, R. Learning dynamical systems from data: An introduction to physics-guided deep learning. Proceedings of the National Academy of Sciences 121, e2311808121 (2024).

32. Lagergren, J. H., Nardini, J. T., Baker, R. E., Simpson, M. J. & Flores, K. B. Biologically-informed neural networks guide mechanistic modeling from sparse experimental data. PLoS Computational Biology 16, e1008462 (2020). 2005.13073.

33. Brunton, S. L., Proctor, J. L. & Kutz, J. N. Discovering governing equations from data by sparse identification of nonlinear dynamical systems. Proceedings of the National Academy of Sciences 113, 3932–3937 (2016). 1509.03580.

34. Rackauckas, C. et al. Universal Differential Equations for Scientific Machine Learning. arXiv (2020). 2001.04385.

35. Philipps, M., Schmid, N. & Hasenauer, J. Universal differential equations for systems biology: Current state and open problems. bioRxiv 2024.11.29.626122 (2024).

36. Fronk, C., Yun, J., Singh, P. & Petzold, L. Bayesian polynomial neural networks and polynomial neural ordinary differential equations. PLOS Computational Biology 20, e1012414 (2024).

37. Daryakenari, N. A., Florio, M. D., Shukla, K. & Karniadakis, G. E. AI-Aristotle: A physics-informed framework for systems biology gray-box identification. PLOS Computational Biology 20, e1011916 (2024).

38. Wang, H. & Hahn, K. M. LOVTRAP: A Versatile Method to Control Protein Function with Light. Current Protocols in Cell Biology 73, 21.10.1–21.10.14 (2016).

39. Kadelka, C., Wheeler, M., Veliz-Cuba, A., Murrugarra, D. & Laubenbacher, R. Modularity of biological systems: a link between structure and function. Journal of the Royal Society Interface 20, 20230505 (2023).

40. Liu, C., Qi, T., Milner, J. J., Lu, Y. & Cao, Y. Speed and Location Both Matter: Antigen Stimulus Dynamics Controls CAR-T Cell Response. Frontiers in Immunology 12, 748768 (2021).

41. Edelstein-Keshet, L. Mathematical Models in Biology 39–71 (2005).

42. Keener, J. & Sneyd, J. Mathematical Physiology, I: Cellular Physiology. Interdisciplinary Applied Mathematics 347–384 (2009).

43. Wang, Y. et al. Mathematical model predicts tumor control patterns induced by fast and slow cytotoxic T lymphocyte killing mechanisms. Scientific Reports 13, 22541 (2023).

44. Rodriguez, J. et al. Predictive nonlinear modeling of malignant myelopoiesis and tyrosine kinase inhibitor therapy. eLife 12, e84149 (2023).

45. McKeithan, T. W. Kinetic proofreading in T-cell receptor signal transduction. Proceedings of the National Academy of Sciences 92, 5042–5046 (1995).

46. Phillips, R., Kondev, J., Theriot, J., Garcia, H. G. & Orme, N. Physical Biology of the Cell 383–425 (2012).

47. Dierckx. Curve and Surface Fitting with Splines. Monographs on numerical analysis (Oxford University Press, 1993). URL https://books.google.com/books?id=-RIQ3SR0sZMC.

48. Mark, G., Gudith, D. & Klocke, U. The cost of interrupted work. Proceedings of the SIGCHI Conference on Human Factors in Computing Systems 107–110 (2008).

49. Khammash, M. H. Perfect adaptation in biology. Cell Systems 12, 509–521 (2021).

50. Li, Y. & Goldbeter, A. Frequency specificity in intercellular communication. Influence of patterns of periodic signaling on target cell responsiveness. Biophysical Journal 55, 125–145 (1989).

51. Li, Y. & Goldbeter, A. Pulsatile signaling in intercellular communication. Periodic stimuli are more efficient than random or chaotic signals in a model based on receptor desensitization. Biophysical Journal 61, 161–171 (1992).

52. Altrock, P. M., Liu, L. L. & Michor, F. The mathematics of cancer: integrating quantitative models. Nature Reviews Cancer 15, 730–745 (2015).

53. Secrier, M., Toni, T. & Stumpf, M. P. H. The ABC of reverse engineering biological signalling systems. Molecular BioSystems 5, 1925–1935 (2009).

54. Toni, T., Welch, D., Strelkowa, N., Ipsen, A. & Stumpf, M. P. Approximate Bayesian computation scheme for parameter inference and model selection in dynamical systems. Journal of The Royal Society Interface 6, 187–202 (2009). 0901.1925.

55. Finkle, J. D., Wu, J. J. & Bagheri, N. Windowed Granger causal inference strategy improves discovery of gene regulatory networks. Proceedings of the National Academy of Sciences 115, 2252–2257 (2018).

56. Rowan, T. The subplex method for unconstrained optimization. PhD thesis, PhD thesis, Ph. D. thesis, Department of Computer Sciences, Univ. of Texas (1990).

